# Parallel pheromone, metabolite, and lipid analyses reveal patterns associated with early life transitions and ovary activation in honey bee (*Apis mellifera*) queens

**DOI:** 10.1101/2024.04.19.590367

**Authors:** Alison McAfee, Armando Alcazar Magaña, Leonard J. Foster, Shelley E. Hoover

## Abstract

Eusocial insects exist in a state of reproductive conflict in which workers forgo reproduction in favor of helping relatives, typically queens, rear offspring. The honest signal hypothesis posits that queens emit pheromonal signals that convey information about their fecundity, which workers use to make decisions around investing in direct vs. indirect fitness and queen acceptance. We evaluated this idea using liquid chromatography-tandem mass spectrometry to measure honey bee queen retinue pheromone (QRP) components in relation to queen age, laying status, and likelihood of acceptance using a protocol that enables QRP to be measured concurrently with metabolomic and lipidomic analyses. We found that older mated queens (>1 month) were more readily accepted by colonies than younger queens (10-12 d), regardless of the queen’s prior laying status. This is despite non-laying queens having significantly smaller ovaries at the time of introduction. Older queens produced higher levels of the QRP components 9(R)-HDA, LEA, and HVA compared to younger queens, with HVA also positively correlating with ovary mass. However, these data suggest that ovary mass is not an influential fertility metric for worker decision-making; therefore, the relationship between HVA and ovary mass is merely an honest signal of a non-influential metric. Metabolomic and lipidomic analyses showed that samples cluster strongly according to queen age and mating status, but not ovary mass. These data also reveal some of the first hints of the importance of prostanoids in queen maturation, along with many other physiological changes that occur in the first month of a queen’s life.

**Significance statement:** Insect pheromones have historically been evaluated using gas chromatography-mass spectrometry, a technique that is incompatible with parallel lipidomics and metabolomics inquests. Here, we repurposed an established two-phase extraction protocol and optimized a liquid chromatography-tandem mass spectrometry method to acquire pheromone, metabolite, and lipid data concurrently from a single sample. We applied this technique to interrogate the honest signal hypothesis, which relates queen pheromone profiles to reproductive quality, but the approach is broadly applicable to any question in which simultaneous determination of complex pheromone profiles and lipidomics or metabolomics data is asset. Such applications may help uncover new pheromones and reveal relationships between pheromones, hormones, and physiology in diverse biological systems.

## Introduction

Components of the honey bee (*Apis mellifera*) queen mandibular pheromone (QMP) were the first pheromones identified among social insect queens^1–3^ and, to date, QMP is the most extensively studied pheromone of honey bees. Now known to contain additional molecules not specific to the mandibular gland ― methyl (Z)-octadec-9-enoate, hexadecan-1-ol, and (Z9,Z12,Z15)-octadeca-9,12,15-trienoic acid, or MO, PA, and LEA, respectively ― as well as an additional mandibular component ― (E)-3-(4-hydroxy-3-methoxyphenyl)-prop-2-en-1-ol, or CA ― the complete blend is better described as queen *retinue* pheromone (QRP)^4^, although QMP is still used to refer to the core components originally identified in mandibular glands: 9-oxo-2(E)-decenoic acid, 9-R- and 9-S-hydroxydec-2(E)-enoic acid, 4-hydroxy-3-methoxyphenylethanol, and methyl p-hydroxybenzoate (abbreviated as 9-ODA, 9(R/S)-HDA, HVA, and HOB, respectively). As the name implies, QRP stimulates worker (sterile females) retinue formation around a queen^1,4^, but the blend and its components also have profound effects on overall colony cohesiveness and continuity by suppressing worker reproduction^5,6^, affecting worker temporal polyethism^7^ and lipid metabolism^8^, drone (male) attraction during mating flights^9^, queen supersedure^10^, swarm cohesion^11^, and worker activity levels via dopamine signalling^12^.

Since the description of QMP in 1988^1^ and extended QRP bouquet in 2003^4^, studies evaluating the pheromone components have exclusively relied on gas chromatography mass spectrometry (GC-MS) for detection and quantification with no major methodological advancements. However, most of the QRP components are compatible with liquid chromatography tandem mass spectrometry (LC-MS/MS), potentially offering seamless integration of pheromone analysis with the burgeoning fields of metabolomics and lipidomics. In particular, the simultaneous lipid and metabolite extraction technique described by Chen *et al.*^13^, which extracts compounds using methylated tert-butyl ether, methanol, and water, followed by phase separation into upper (organic, lipid) and lower (aqueous, metabolite) fractions, would allow for parallel pheromone, metabolite, and lipid extraction from a single sample. Along with chromatography methods optimized for separating hydroxylated fatty acids and their isomers, such a technique could provide exceptionally rich data for drawing relationships between pheromone profiles, physiological changes, and individual phenotypes such as queen age or reproductive quality.

Evolutionary theory and positive relationships between QMP and fertility metrics have lent support to what is known as the “honest signal hypothesis,” whereby workers may weigh indirect fitness costs and benefits of supporting the existing queen compared to rearing a new queen, or the direct fitness benefits of producing unfertilized eggs (sons) themselves^14–16^. In this model, queen-derived pheromones act as messages to the workers, which convey information about the queen’s fertility status or reproductive quality, but there is evidence both for and against this scenario. For example, workers are most attracted to naturally mated queens with larger ovaries^17,18^, and queens with larger ovaries produce more QMP^18^. However, these studies did not utilize age-matched queens, and others have argued that since the primary QMP component (9-ODA) does not differ between infertile (drone-laying), and age-matched fertile (worker-laying) queens, QMP is not an honest signal of fertility (though notably, other QMP components – HVA and 9-HDA – positively correlated with fertility in the same study)^19^.

Replacing old or unproductive queens with young, vigorous queens or those from genetically desirable stock is a standard procedure in honey bee colony management^20^. If a queen is rejected, the colony may lose productivity (due to a long broodless period) or perish (if a new queen is not successfully reared). If the honest signal hypothesis is true, one might expect beekeepers to have higher requeening success if queens with the highest reproductive potential are used; however, additional factors contribute to whether a queen is perceived as “desirable” by workers. For instance, Rhodes *et al.*^21^ found that queen acceptance rates increase with age (within a range of 7-35 d post-emergence). Additionally, worker sensitivity to QMP varies throughout the season^22–24^. Likelihood of acceptance also appears to depend on the honey bee subspecies in question^24^ and whether the queen and recipient colony belong to matching subspecies^25^.

This wide range of influential factors may explain why introducing mated queens to new colonies can have such variable success, with as few as 45% or as high as 95% of colonies accepting their new queen using the standard delayed release method^21,22,26,27^, in which workers are allowed to acclimate to their new queen while she is constrained and protected in a small cage for several days before being freed. This can dramatically impact the colony’s likelihood of survival; if queen rejection goes undetected by the beekeeper, workers must then either rear a new queen or become hopelessly queenless, ultimately perishing. Moreover, colonies experiencing “queen events” (queen loss or evidence of queen replacement) are over three times more likely to die over the following 50 days compared to colonies not experiencing queen events^28^, in part due to the riskiness of requeening.

Understanding the influence of factors that can be controlled during the queen production and distribution process, such as queen age or banking, on queen acceptance and physiology is therefore of both practical importance to beekeepers and biological significance under the framework of the honest signal hypothesis. Mated queens are routinely banked (temporarily held in small cages within larger colonies) ahead of distribution to recipient beekeepers; however, the impact this may have on subsequent acceptance rates is not known. This scenario is also particularly relevant for investigating the honest signal hypothesis, since banking offers a method of manipulating (restricting) queen laying and perhaps ovary mass while keeping other queen variables (*e.g.* source, age, mating quality, *etc*.) constant or randomly distributed.

Here, we re-examined the influence of queen age and pheromone profiles on acceptance rates as originally reported by Rhodes *et al.*^21,29^, while also comparing acceptance rates, metabolites, lipids, and pheromone profiles of banked queens to age-matched free-range queens. This design allowed us to differentiate between the influence of age and ovary activation, providing a new understanding of the honest signal hypothesis. We evaluated queen pheromones, metabolites, and lipids using a two-phase extraction based on a previously published approach by Chen *et al.*^13^, which capitalizes on the high-sensitivity and rich data generated by ultra-high performance LC-MS/MS instead of GC-MS, which has historically been the technique of choice for QRP analysis. To do this, we developed a new UHPLC method for deep coverage of polar lipids, including the resolution of structural isomers of hydroxylated short chain fatty acids such as 10- and 9-hydroxydec-2(E)-enoic acid. This approach is compatible with parallel metabolomics and lipidomics data acquisition, providing rich information on covarying hormones and substrates.

## Methods

### Queen acceptance rates

A schematic of our experimental design is presented in **Figure 1**. Queens were produced by Scandia Honey in May 2023 in Scandia, AB, Canada, according to standard queen rearing methods. Briefly, very young (≤ 1 day old) larvae were grafted from a single mother colony into queen cell cups (JZBZ) and reared into queen cells in strong cell builder colonies. Ten days after grafting (June 1), queen cells were placed in small nucleus colonies (nucs) to facilitate mating. The expected emergence dates based on average queen developmental times were between June 3^rd^ and 4^th^. On June 5^th^, N = 10 newly emerged queens (the “virgin” group) were sampled in JZBZ plastic cages and transported to the laboratory (Lethbridge, AB) in a battery box with attendant nurses, where they were euthanized, frozen at -80 °C, then tipped into 1.5 mL microfuge tubes and stored at -80 °C.

**Figure 1.**
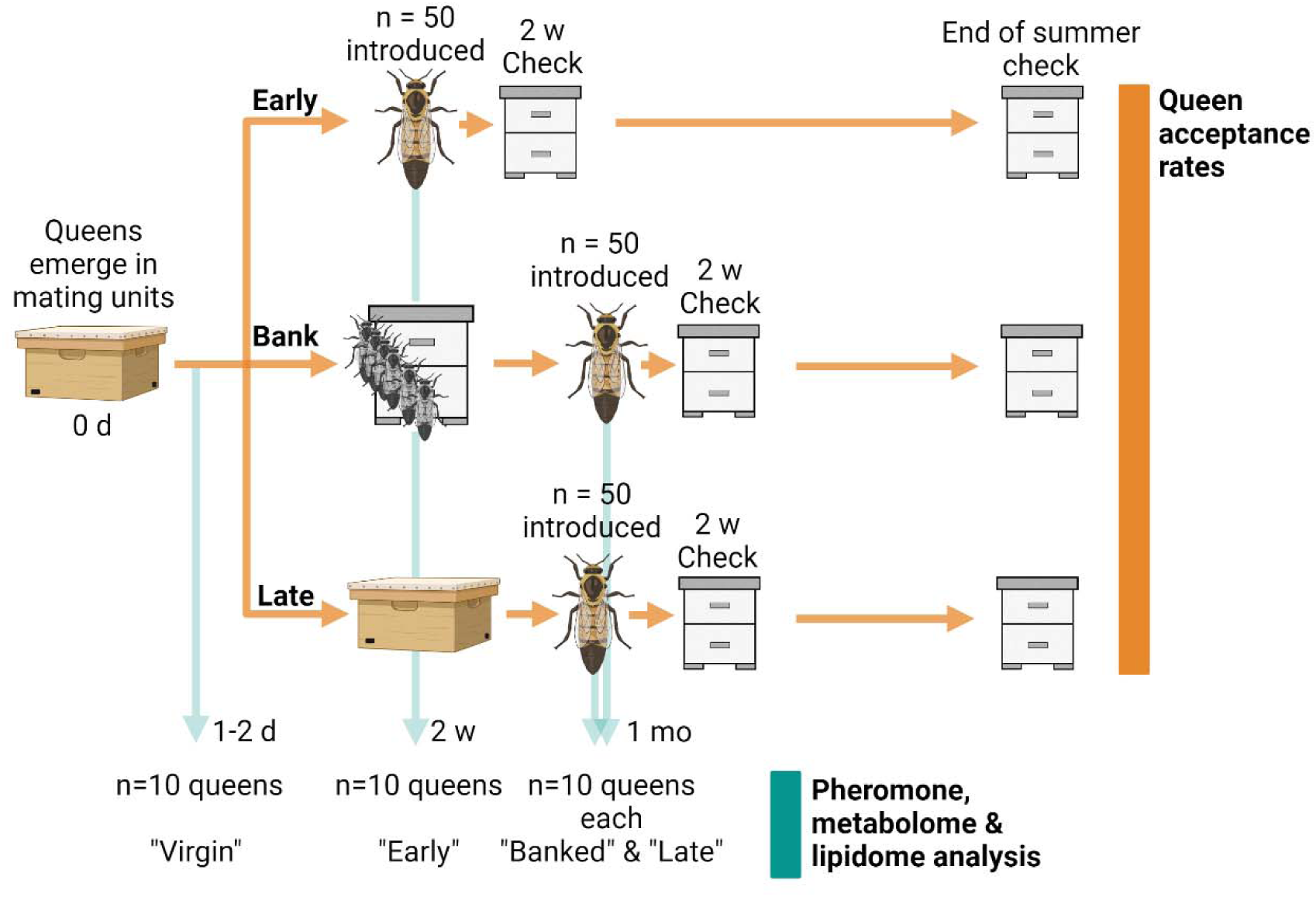
Schematic of the field experimental design. Queens were reared simultaneously and separated into three cohorts: Early (sampled and introduced to new colonies 10-12 d after emergence), banked (sampled and introduced to queen banks for 18 days, then introduced to new colonies 1 month after emergence), and late (aged free-range in mating nucs, then introduced to new colonies 1 month after emergence) cohorts. See methods for complete experimental details. Figure created with Biorender.com.

Two weeks after queen cells were placed in mating nucs (June 15), young queens belonging to the “early” group were sampled, with one cohort euthanized for molecular analysis, a second cohort introduced into queenless colonies to evaluate acceptance rates, and a third cohort moved into three separate queen bank colonies. N = 10 mated queens (confirmed to be laying eggs in the mating nucs) were sampled and stored in the same fashion as the queens above for metabolomics and lipidomics analysis, whereas another N = 50 early mated queens were caged and introduced via a 2.5 cm candy tube into double brood chamber recipient colonies which had been de-queened the previous day. Introduction success (judged by the marked queen being present and laying worker brood) of the early mated queens was evaluated on June 28 and sustained acceptance was recorded on September 30. N = 70 young, mated queens were moved into queen banks to eventually serve as the “banked” group. The remaining mated queens were allowed to mature (“free-range”) in their respective mating nucs until the next sampling event.

Approximately one month after queen emergence (July 4), queens belonging to the late and banked groups were sampled, again with a subset retained for molecular analysis (N = 10 each) and a larger cohort introduced to colonies (N = 50 each) as described above. Two weeks after introduction, late and banked recipient colonies were checked for queen acceptance, and colonies were checked again on September 30.

### Pheromone standards

Chemical standards for queen retinue pheromone components were obtained from the sources listed in **Table 1**. We included all core QRP components, as well as 10-hydroxy-2(E)-decanoic acid (10-HDA), which is chemically similar to 9-HDA and is also produced in the mandibular glands. 10-HDA is considered to be a worker-like pheromone tending to be elevated in virgin queens^30^ (although there is disagreement in the literature^31^), and is a major component of royal jelly^32^. Standards were dissolved in methanol and serially diluted in 80% methanol (with 0.01% butylated hydroxytoluene, or BHT) to a final concentration of 1 ppm. Standards were used for method optimization and were analyzed at the beginning and end of the sample batch under identical conditions to determine retention times, isotopic distributions, and fragmentation patterns. This enabled the compound peaks to be confidently assigned and extracted from the data acquired from untargeted analysis.

**Table 1.** Pheromone standards, abbreviations, and suppliers.

| Compound | Abbr. | Monoisotopic mass (Da) | Source |
| --- | --- | --- | --- |
| 9-oxo-2(E)-decenoic acid | 9-ODA | 184.1099 | Intko Supply Ltd,<br>Chilliwack, BC, Canada |
| 9-hydroxy-2(E)-decenoic acid* | 9-HDA | 186.1256 | Intko Supply Ltd,<br>Chilliwack, BC, Canada |
| methyl p-hydroxybenzoate | HOB | 152.0473 | MilliporeSigma,<br>Burlington, MA, USA |
| 4-hydroxy-3-methoxyphenylethanol | HVA | 168.0786 | MilliporeSigma,<br>Burlington, MA, USA |
| methyl (Z)-octadec-9-enoate | MO | 296.2715 | Cayman Chemical, Ann Arbor, MI, USA |
| E-3-(4-hydroxy-3-methoxyphenyl)-prop-2-en-1-ol | CA | 180.0786 | Cayman Chemical, Ann Arbor, MI, USA |
| hexadecan-1-ol | PA | 242.2610 | VWR International,<br>Radnor, PA, USA |
| (Z9,Z12,Z15)-octadeca-9,12,15-trienoic acid | LEA | 278.2246 | Cayman Chemical, Ann Arbor, MI, USA |
| 10-Hydroxy-2(E)-decenoic acid | 10-HDA | 186.1256 | Cayman Chemical, Ann Arbor, MI, USA |

\*R and S 9-HDA enantiomers were only commercially available in a mixed form (85% R, 15% S). In the supplied mixture, only the R enantiomer was observable; therefore, all 9-HDA relative quantities in this manuscript refer to the R enantiomer (although the S enantiomer does appear to be identifiable in samples; **Figure S1**).

### Metabolomics and lipidomics sample processing

Untargeted metabolomics and lipidomics analyses were conducted on queen head extracts in order to perform relative quantification of retinue pheromone components, metabolites, and lipids (**Figure 2**). A step-by-step protocol is included in **Supplementary File 1.** The virgin, early, late, and banked queens (40 samples total) which were retained for molecular analysis were shipped to UBC on dry ice and stored at -70 °C until processing. To minimize matrix effects and achieve low detection limits, a two-phase extraction was conducted according to methods previously described^13,33^. For all queens, ovaries were dissected and weighed. Then, whole heads were dissected and deposited into homogenization tubes, along with 4 ceramic beads and 400 µl chilled (-20 °C) extraction solvent (75% methanol (v/v), 25% water (v/v), and 0.01% BHT (w/v)). BHT inhibits the oxidation or isomerization of reactive compounds throughout the extraction process, reducing the likelihood of potential artifacts. Three “blank” samples were processed in parallel, consisting of a homogenization tube with clean ceramic beads, following the same steps as for the samples. In order to track the performance of the method, the extraction solvent was spiked with the internal standards methionine-d3, ferulic acid-d3, caffeine-^13^C_3_, and CUDA (12-[[(cyclohexylamino)carbonyl]amino]-dodecanoic acid at 1 ppm, and 5 µl of SPLASH® LIPIDOMIX® mass spec standard (Avanti Polar Lipids, Birmingham, AL, USA) containing a deuterated mixture of 14 major lipid classes. These internal standards facilitate identifying the elution region of each lipid class within the chromatogram. Samples were homogenized using a Precellys 24 tissue homogenizer (3 x 30 s at 5,000 s^-1^, 2 min rest on ice in between), then 1 mL of chilled (-20 °C) methyl tert-butyl ether (MTBE) was added. Samples were vortexed, then incubated for 1 hour (20 °C, shaking at 1,000 rpm) for two-phase extraction. After this incubation, debris was pelleted (14,000 *g*, 10 min, 4 °C) and 1.2 mL of supernatant was transferred to a new tube. Phase separation was induced by adding 214 µl ddH_2_O (to a final additional concentration of 15%), after which the samples were vortexed and incubated at room temperature for 10 min. Finally, samples were centrifuged (14,000 *g*, 15 min, 4 °C) and a 250 µl aliquot was extracted from both the upper phase (containing less polar compounds, mainly lipids), and the lower phase (containing more polar compounds, mainly metabolites). The metabolite samples were dried in a speedvac (2.5 h, room temperature) and stored at -70 °C, whereas the lipid samples were stored at -70 °C in solution, then dried in a speedvac immediately ahead of analysis.

**Figure 2.**
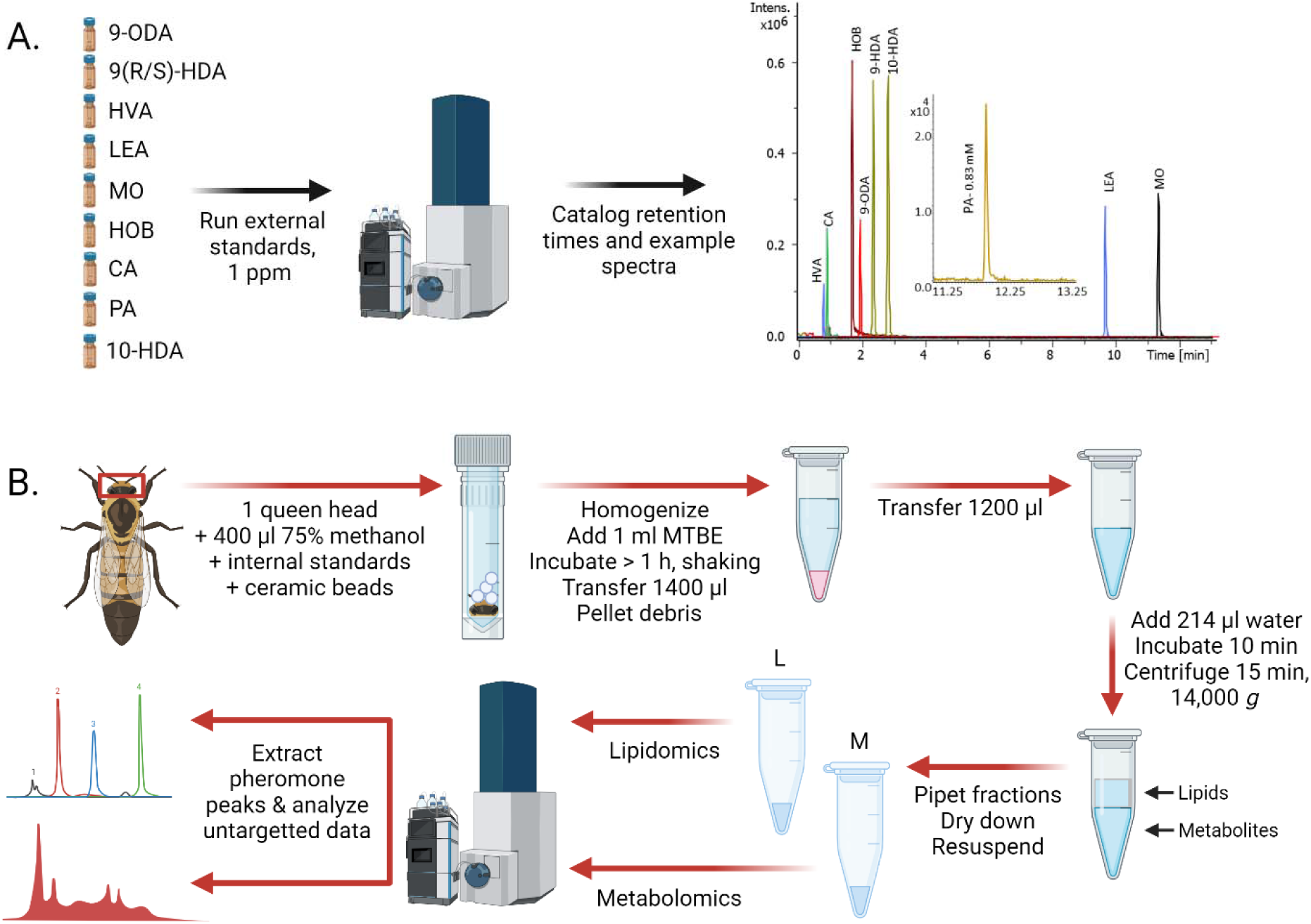
Schematic of the analytical workflow. The extraction protocol is adapted from Chen *et al.*^13^ A) External pheromone standards are first analyzed to determine retention times and spectra to confirm peak assignments in subsequent untargeted data analysis. B) Then, metabolomics and lipidomics samples are extracted from whole queen heads. Pheromone components tend to partition in the organic (lipid) phase but HOB and CA were also measured in the aqueous (metabolite) phase. This approach allows for relative quantification of pheromones concurrently with thousands of other lipid and metabolite features. Figure created with Biorender.com.

### Metabolomics LC-MS/MS analysis

The polar fraction was resuspended in 250 µl of 50% aqueous methanol (v/v). The suspension was centrifuged at 16,000 rcf for 10 min, and 200 µl of the resulting supernatant was transferred to LC-MS vials (Thermo Fisher) for LC-MS/MS analysis. Aliquots of 20 µL from each sample were pooled to generate a quality control sample (QC) used for evaluating analytical performance. Quality control samples and blanks were injected every 10 samples. To avoid artifacts due to batch effects, samples were fully randomized before injecting 3 µl and 2 µL in positive and negative ionization mode, respectively. Extracted compounds were analyzed using an Impact^TM^ II high-resolution mass spectrometer (Bruker Daltonics, Bremen, Germany) coupled with a Vanquish Horizon UHPLC system (Thermo). Separation of compounds was achieved using a multigradient method on an Inertsil Ph-3 UHPLC column (2 µm, 150 x 2.1 mm) (GL Sciences) equipped with a Ph-3 guard column (2 µm, 2.1 x 10 mm). The mobile phase consisted of water (A) supplemented with 0.1% (v/v) formic acid, and methanol (B) with 0.1% (v/v) formic acid. The chromatographic separation utilized a multi-step gradient from 5% to 99% mobile phase B over 18 min. The UHPLC program was set as follows: 0 min (5% B), 0–1 min (5% B), 1–8 min (35% B), 8–10.5 min (99% B), 10.5–14 min (99% B), 14–14.5 min (5% B), and 14.5–18 min (5% B). The column temperature was set to 55 °C, while the autosampler was maintained at 4 °C, and the flow rate was 0.3 mL/min. Data-dependent acquisitions were conducted in negative (ESI-) and positive (ESI+) ionization modes to obtain precursor and fragment ion information for annotating compounds. For positive ion mode, the mass spectrometer settings were as follows: capillary voltage of 4,500 V, nebulizer gas pressure of 2.0 bar, dry gas flow rate of 9 L/min, dry gas temperature of 220 °C, mass scan range of 60-1,300 m/z, and a total cycle time of 0.6 s. To obtain comprehensive structural information, collision energy of 20 V was ramped through each MS/MS scan from 100 to 250%. For negative ionization mode, the capillary voltage was set at -3,500 V. To ensure high mass accuracy, internal calibration was conducted in each analytical run using 10 µl of 10 mM sodium formate injected at the start of the run (from 0 to 0.15 min) via a 6-port valve (average mass error was below 1.5 ppm, **Supplementary Data 1-4**).

### Lipidomics LC-MS/MS analysis

We were particularly interested in resolving 9- and 10-HDA as well as several prostanoids because of their distinct roles in queen bees and other insects^31,34^. To accomplish this, we tested various gradients of water, methanol, acetonitrile, and isopropanol. A critical approach to identifying and resolving numerous polar lipids was to determine the appropriate polarity at the start of chromatography to retain polar lipids without precipitation at the beginning of the injection. Excessive water causes lipid precipitation, resulting in broad peaks with low sensitivity. Similarly, excessive organic solvent elutes polar lipids near the void volume of the column. In traditional lipidomics methods^35^, polar lipids such as hydroxylated short-chain fatty acids, prostaglandins and thromboxanes all elute in the first two minutes of the chromatogram, leading to poor resolution of isomers and considerable peak suppression. For instance, the internal standard CUDA elutes roughly at 0.75 minutes in traditional settings^35^. However, in our setting, this internal standard elutes at minute 7.0. Our final optimized methods are detailed below.

Dried lipid extracts were resuspended in 250 µl of a solution containing 70% acetonitrile and 30% isopropanol (v/v) on the day of the LC–MS/MS analysis as described in the metabolomics section. Lipids were separated on an ACQUITY UPLC CSH C18 analytical column (130Å, 1.7 µm, 2.1 mm X 100 mm, Waters) with a multi-step elution gradient optimized to resolve polar lipids such as hydroxylated fatty acids. The mobile phase consists of water (A) supplemented water with 10 mM ammonium formate and 0.1% formic acid, and (B) 10% acetonitrile and 90% isopropanol (v/v) supplemented with 10 mM ammonium formate and 0.1% (v/v) formic acid. The chromatographic separation utilized a multi-step gradient from 20% to 99% mobile phase B over 18 min. The UHPLC program was set as follows: 0 min (20% B), 0–2 min (20% B), 2–11 min (80% B), 11–11.5 min (99% B), 11.5–13.2 min (99% B), 13.2–14 min (5% B), and 14–18 min (5% B). The column was operating at 0.4 mL/min and 65 °C. Sample temperature was maintained at 4 °C. One and 2 µl were injected for positive and negative ionization mode, respectively. Data were collected using data-dependent high-resolution mass spectral acquisition (Bruker Impact II) in ESI+ and ESI-. For ESI+, the mass spectrometer settings were as follows: capillary voltage of 4,500 V, nebulizer gas pressure of 2.0 bar, dry gas flow rate of 9 L/min, dry gas temperature of 220 °C, mass scan range of 100-1,700 m/z. To achieve a lower limit of detection, the spectra acquisition rate was set at 3 Hz and a cycle time of 0.6 s. The collision energy of 20 V was ramped through each MS/MS scan from 100 to 250%. For ESI-, the capillary voltage was set at -3,800 V.

### Metabolomics and lipidomics data processing

Raw data processing was conducted using Progenesis QI™ software (V3.0.7600.27622) with the METLIN^TM^ plugin V1.0.7642.33805 (NonLinear Dynamics), encompassing peak picking, alignment, deconvolution, normalization, and database querying and searching, as described previously^36,37^. To increase annotation confidence, deconvoluted ions were considered for annotation only if they fulfilled the following criteria: 1) compounds contained structural information (MS/MS), 2) the coefficient of variation (CV) was below 25% across all QC samples, 3) intensity in samples was >5-fold the intensity in experimental blanks, and 4) the mass error was below 5 ppm (average mass error in annotation below 1.5 ppm, **Supplementary Data 1-4**). Candidate annotation confidence was in accordance with reporting criteria for chemical analysis suggested by the Metabolomics Standards Initiative (MSI) guided by a Progenesis QI score^38,39^. Level one (L1) annotations were acquired through matching against the Life Science Institute (UBC) *in-house* spectral library (Mass Spectrometry Metabolite Library of Standards, MSMLS, supplied by IROA Technologies) containing over 500 standards with broad representation of primary metabolism and injected under identical conditions. For level two (L2) annotations, features were screened against METLIN^TM^, GNPS, HMDB, and MassBank of North America^40–43^. LipidBlast^44^ was also included for lipidomics analysis. When compounds were detected in both ion modes, the one with a lower CV in QCs was retained. Relative metabolite quantities were computed through peak area calculations, with normalization conducted using "all compounds normalization", a robust Progenesis QI^TM^ built-in approach designed explicitly for untargeted metabolomics (www.nonlinear.com).

### Statistical analysis

All statistical analyses were performed in R (version 4.3.0) using R Studio (version 2023.09.1+494)^45^. Packages utilized include those within the tidyverse^46^ (ggplot2, dplyr, tibble, and readxl), as well as ggfortify^47^, reshape^48^, limma^49^, DHARMa^50^, car^51^, and emmeans^52^.

For the queen acceptance data, statistical comparisons between groups were made using a logistic regression, which modelled queen acceptance (1 = accepted, 0 = rejected) by group (early, late, and banked) at each time point (2 weeks after introduction or end of summer) separately. Appropriateness of fit was determined by using the DHARMa package to inspect simulated residual distributions^50^. Ovary masses of sampled queens were compared using a linear model, with *post hoc* contrast statistics extracted using emmeans^52^.

For the pheromone data, peak areas for each compound were integrated using Progenesis QI™ (see **Figure S1-S3** for example chromatograms) and log_10_-transformed, then differences between groups were analysed using linear models, except in the case of 9-ODA, which was analyzed using a Kruskal-Wallis test owing to unequal variance between groups and resultant skewed residual distributions. The linear model included ovary mass and queen group (four levels) as an interactive term, but if no significant effect of ovary mass was observed, the term was dropped from the model. As for the queen acceptance data, residual distributions resulting from the linear models were inspected for appropriateness of fit. *Post hoc* contrast statistics were extracted using emmeans^52^.

For the metabolomics and lipidomics data, raw intensity data were first log_10_-transformed and the global distributions were inspected for normality. Principal component analyses (PCAs) were conducted on each data set separately, combining all compounds identified in both positive and negative mode. Preliminary testing was conducted on data from each ion mode separately, as well as with the full list of features compared to a reduced list of only features with annotations, showed that clustering patterns were similar in all cases. Therefore, all features and both ion modes were used to generate the final PCA plots using the prcomp and autoplot functions in R. Variable importance in projection (VIP) scores were calculated using MetaboAnalyst 5.0^53^.

Global differential abundance analyses were performed using the limma package^49^ on metabolomics and lipidomics data separately. Family-wise false-discovery rates for each contrast were controlled to 1% using the Benjamini-Hochberg method. Scaled (z-scored) data were clustered using the hclust function and plotted in the order of clustering using ggplot2^54^. In addition, among the mated queens, relationships between ovary mass and compound abundance were evaluated following the same methods as described above, except that ovary mass was used as the dependent variable.

### Data availability

All of the data and code underlying the analyses we present are publicly available. R codes are provided as **Supplementary File 1**, processed metabolomics and lipidomics data (along with statistical results and annotation confidence) are provided as **Supplementary Data 1-4**, and raw mass spectrometry data are available at Metabolomics Workbench (www.metabolomicsworkbench.org; Accession: PR001924; doi: http://dx.doi.org/10.21228/M81B11)

## Results

### Queen acceptance rates

Queen acceptance likelihood did not significantly depend on group when measured two weeks after introduction (logistic regression; Type II Wald χ^2^ = 2.5, 3 levels, N = 150, p = 0.29; **Figure 3A**) or at the end of summer (approximately three months after introduction; logistic regression; Type II Wald χ^2^ = 3.8, 3 levels, N = 150, p = 0.15; **Figure 3B**). However, the “early” queens (introduced to new colonies < 2 weeks after emergence) tended to have the lowest acceptance rates, with the most pronounced difference occurring at the end-of-summer time point. The biggest difference was between early and banked cohort queens, with the banked cohort having 14% better acceptance rates than the early cohort (90% vs. 76%; logistic regression; z = 1.8, p = 0.070). The trend of banked queens having high acceptance rates is not positively linked to the queen’s ovary mass at introduction, as banked queens had significantly smaller ovaries than early cohort queens (linear model; Tukey adjustment method, t = 3.2, df = 27, p = 0.0090; 27% smaller) and late cohort queens (t = 2.8, p = 0.026; 24% smaller) (**Figure 3C**).

**Figure 3.**
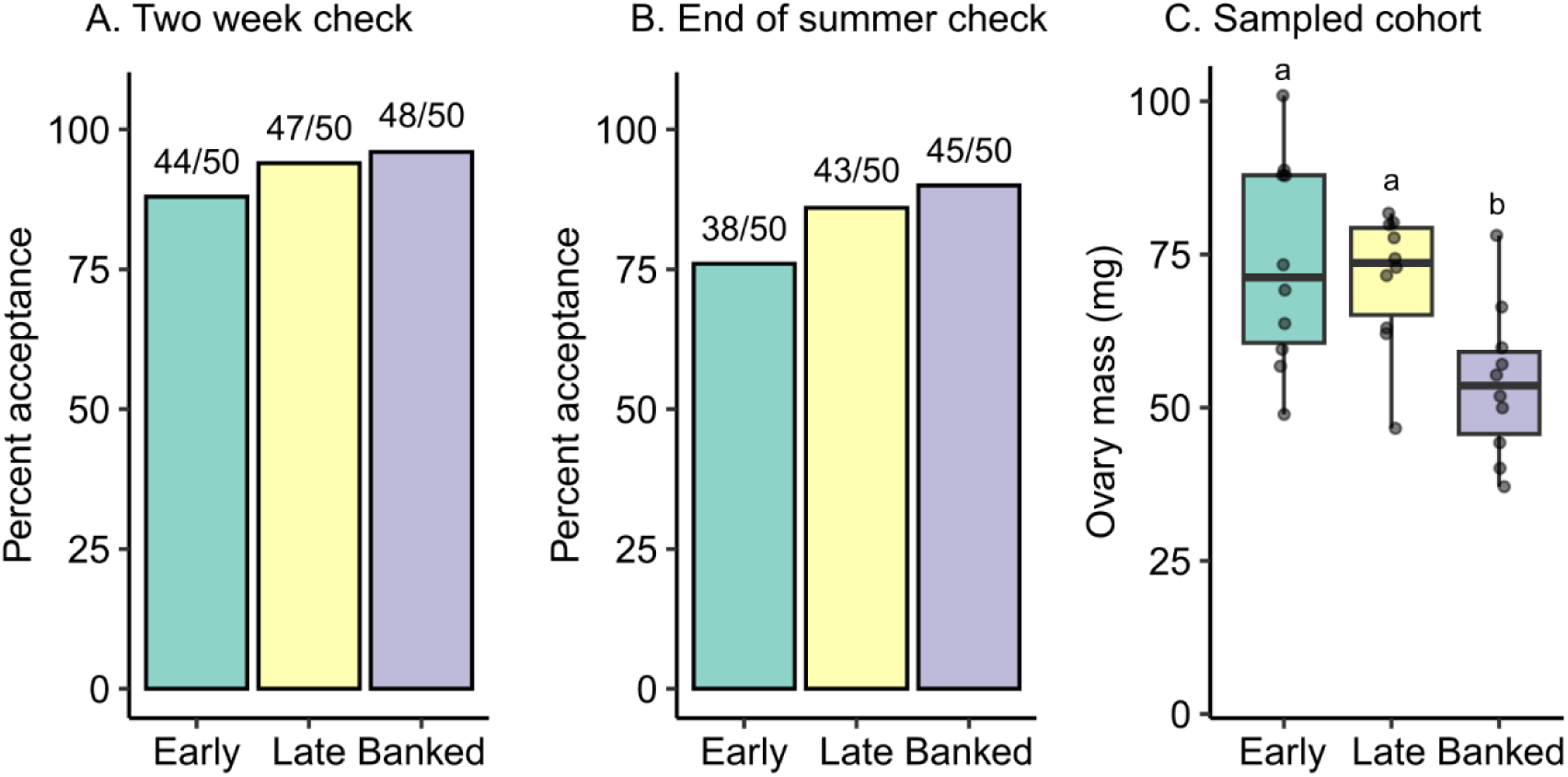
Ovary mass at introduction is not significantly linked to acceptance likelihood. Queens from mating nucs were introduced into new colonies 10-12 days after emergence (early), one month after emergence (late), or removed from mating nucs 10-12 days after emergence then banked for 18 days before being introduced to new colonies (banked). A) Queen acceptance rates two weeks after introduction. There is no significant relationship with group B) On September 30, queen acceptance was recorded for all groups (3.5 months post-introduction for early queens, 3 months post-introduction for late and banked queens). There is no significant main effect of group, but banked queens tend to have higher acceptance rates than early queens (logistic regression, z = 1.8, p = 0.070). C) A cohort of 10 queens from each group were reserved on each introduction day for laboratory analysis. Among these queens, ovary mass varied by group, driven by the small ovaries of banked queens (linear model; df = 27, F = 6.1, p = 0.0065).

### Queen retinue pheromone profiles

Analysis of 10-HDA, 9-ODA, 9-HDA, HOB, HVA, LEA, MO, and CA (example chromatograms in **Figure S1-S3**) revealed that all pheromones except MO show significant differences between groups, with virgin queens often producing relatively low amounts, except for 10-HDA, which was present in the highest amounts in virgins (**Figure 4A**). The virgin group was not among our queen introduction cohorts, but was included here in part to compare our ability to detect differences using LC-MS/MS, which has not been previously used to measure queen pheromones, to existing literature comparing virgin and mated queens^31^. Among the mated queens, older queens (banked and/or late cohorts) produced lower levels of 10-HDA and CA than younger queens (the early cohort), but higher levels of 9-HDA, HVA, and LEA. In addition, HVA significantly correlated with ovary mass (**Figure 4B**) and 9-HDA varied according to a significant interaction between ovary mass and cohort (**Figure 4C**). No pheromones were significantly different between banked and age-matched free-range (late) queens after regressing out the influence of ovary mass. Summary statistics for all pheromone comparisons are in **Table 2**. Long-chain alcohols often have lower ionization efficiency; consequently, the pheromone hexadecane-1-ol (PA) was below our limit of detection (experimentally determined to be 79 µM on our system). Example MS/MS spectra for QRP components in standards and samples are shown in **Figure S4**.

**Figure 4.**
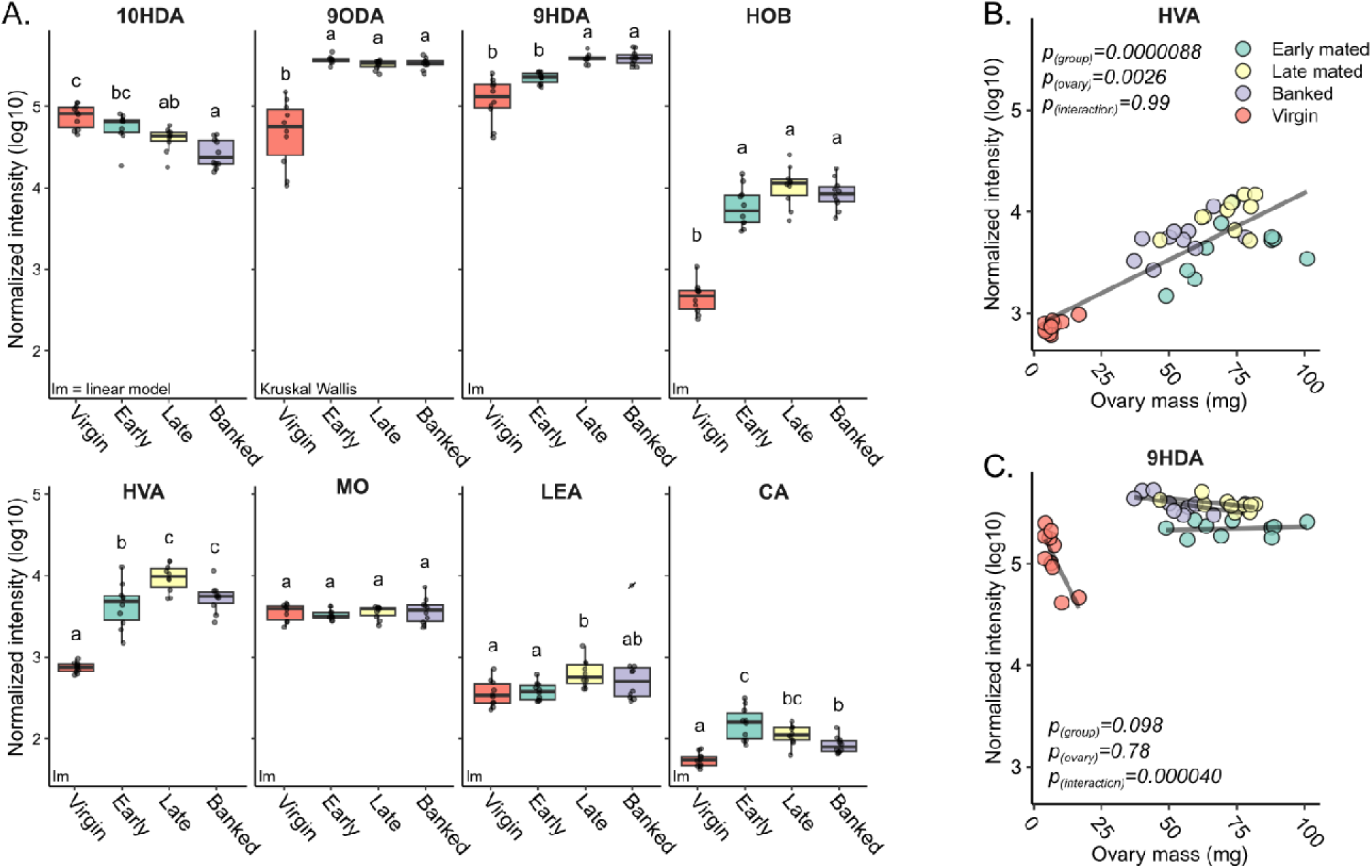
Queen retinue pheromone components vary by group and ovary mass. We performed two-phase extraction to obtain queen pheromone components from whole heads. A) Log_10_-transformed, normalized pheromone component peak areas were analyzed using a linear model, including group and ovary mass as interacting factors, except for 9-ODA. If no significant relationships with ovary mass were identified, the term was dropped from the model. Due to unequal variance, 9-ODA variation by group was analyzed using a Kruskal-Wallis test. Letters above boxes indicate significantly different groups. HVA (B) and 9-HDA (C) abundance covaried with ovary mass as main effect and interactive effect, respectively. 10HDA = 10-hydroxy-2(E)-decanoic acid, 9ODA = E-9-oxodec-2-enoic acid, 9HDA = 9(R)-hydroxydec-2(E)-enoic acid, HOB = methyl p-hydroxybenzoate, HVA = 4-hydroxy-3-methoxyphenylethanol, MO = methyl oleate, LEA = linolenic acid, CA = E-3-(4-hydroxy-3-methoxyphenyl)-prop-2-en-1-ol. All assignments were confirmed against synthetic standards. Summary statistics for A-C are shown in **Table 2**.

**Table 2.**
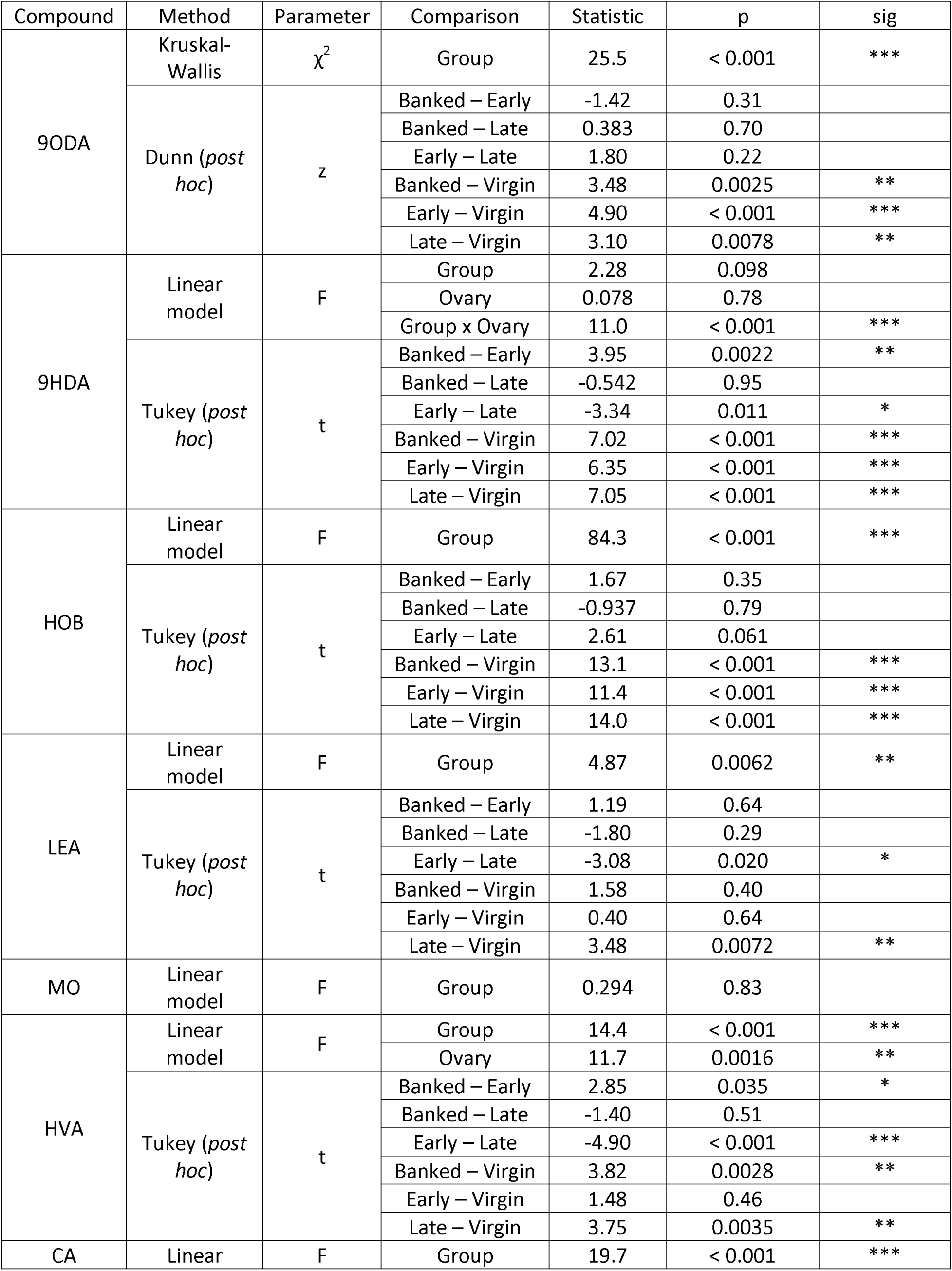
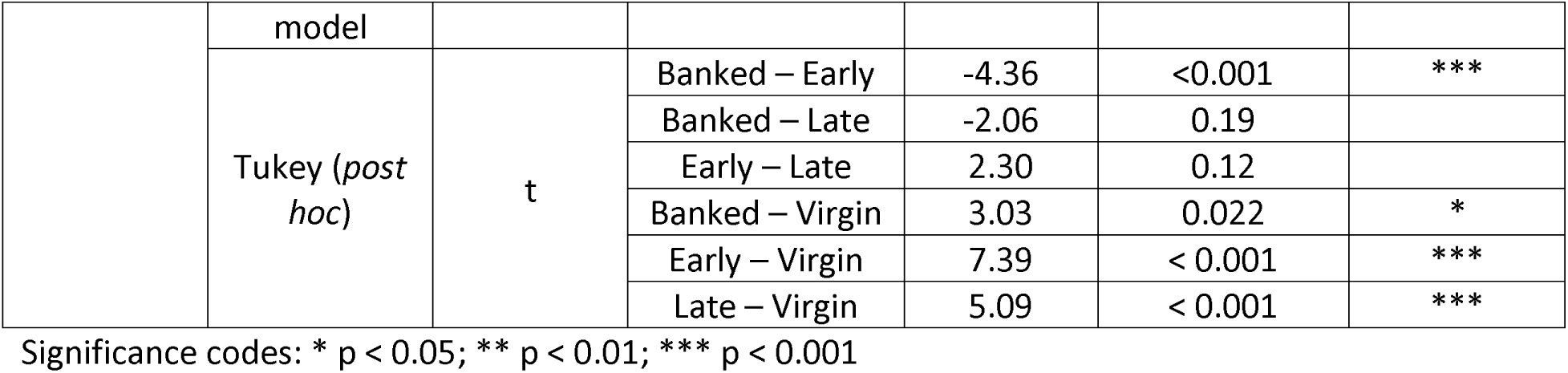
Summary statistics for pheromone comparisons.

| Compound | Method | Parameter | Comparison | Statistic | p | sig |
| --- | --- | --- | --- | --- | --- | --- |
| 9ODA | Kruskal-Wallis | $\chi^2$ | Group | 25.5 | < 0.001 | *** |
|  | Dunn (*post hoc*) | z | Banked – Early | -1.42 | 0.31 |  |
|  |  |  | Banked – Late | 0.383 | 0.70 |  |
|  |  |  | Early – Late | 1.80 | 0.22 |  |
|  |  |  | Banked – Virgin | 3.48 | 0.0025 | ** |
|  |  |  | Early – Virgin | 4.90 | < 0.001 | *** |
|  |  |  | Late – Virgin | 3.10 | 0.0078 | ** |
| 9HDA | Linear model | F | Group | 2.28 | 0.098 |  |
|  |  |  | Ovary | 0.078 | 0.78 |  |
|  |  |  | Group x Ovary | 11.0 | < 0.001 | *** |
|  | Tukey (*post hoc*) | t | Banked – Early | 3.95 | 0.0022 | ** |
|  |  |  | Banked – Late | -0.542 | 0.95 |  |
|  |  |  | Early – Late | -3.34 | 0.011 | * |
|  |  |  | Banked – Virgin | 7.02 | < 0.001 | *** |
|  |  |  | Early – Virgin | 6.35 | < 0.001 | *** |
|  |  |  | Late – Virgin | 7.05 | < 0.001 | *** |
| HOB | Linear model | F | Group | 84.3 | < 0.001 | *** |
|  | Tukey (*post hoc*) | t | Banked – Early | 1.67 | 0.35 |  |
|  |  |  | Banked – Late | -0.937 | 0.79 |  |
|  |  |  | Early – Late | 2.61 | 0.061 |  |
|  |  |  | Banked – Virgin | 13.1 | < 0.001 | *** |
|  |  |  | Early – Virgin | 11.4 | < 0.001 | *** |
|  |  |  | Late – Virgin | 14.0 | < 0.001 | *** |
| LEA | Linear model | F | Group | 4.87 | 0.0062 | ** |
|  | Tukey (*post hoc*) | t | Banked – Early | 1.19 | 0.64 |  |
|  |  |  | Banked – Late | -1.80 | 0.29 |  |
|  |  |  | Early – Late | -3.08 | 0.020 | * |
|  |  |  | Banked – Virgin | 1.58 | 0.40 |  |
|  |  |  | Early – Virgin | 0.40 | 0.64 |  |
|  |  |  | Late – Virgin | 3.48 | 0.0072 | ** |
| MO | Linear model | F | Group | 0.294 | 0.83 |  |
| HVA | Linear model | F | Group | 14.4 | < 0.001 | *** |
|  |  |  | Ovary | 11.7 | 0.0016 | ** |
|  | Tukey (*post hoc*) | t | Banked – Early | 2.85 | 0.035 | * |
|  |  |  | Banked – Late | -1.40 | 0.51 |  |
|  |  |  | Early – Late | -4.90 | < 0.001 | *** |
|  |  |  | Banked – Virgin | 3.82 | 0.0028 | ** |
|  |  |  | Early – Virgin | 1.48 | 0.46 |  |
|  |  |  | Late – Virgin | 3.75 | 0.0035 | ** |
| CA | Linear | F | Group | 19.7 | < 0.001 | *** |
|  | model |  |  |  |  |  |
|  | Tukey (*post hoc*) | t | Banked – Early | -4.36 | <0.001 | *** |
|  |  |  | Banked – Late | -2.06 | 0.19 |  |
|  |  |  | Early – Late | 2.30 | 0.12 |  |
|  |  |  | Banked – Virgin | 3.03 | 0.022 | * |
|  |  |  | Early – Virgin | 7.39 | < 0.001 | *** |
|  |  |  | Late – Virgin | 5.09 | < 0.001 | *** |

Significance codes: \* $p < 0.05$ ; \*\* $p < 0.01$ ; \*\*\* $p < 0.001$

### Metabolomics and lipidomics

In the metabolite analysis, we detected 1,346 unique molecular features in positive ion mode and 1,677 in negative ion mode, 100 and 138 of which were annotated, respectively, according to the Metabolomics Standard Initiative (MSI) criteria (detailed scores and annotation level in **Supplemental Data 1-4**)^38^. In the lipid analysis, we detected 1,111 unique molecular features in positive ion mode and 628 in negative ion mode, 112 and 146 of which were annotated. Principal component analysis of metabolomics and lipidomics fractions show that samples cluster strongly according to queen age and not banking status (**Figure 5**), with the combined principal components accounting for 58.4% and 58.1% of the variation in metabolomics and lipidomics data, respectively.

**Figure 5.**
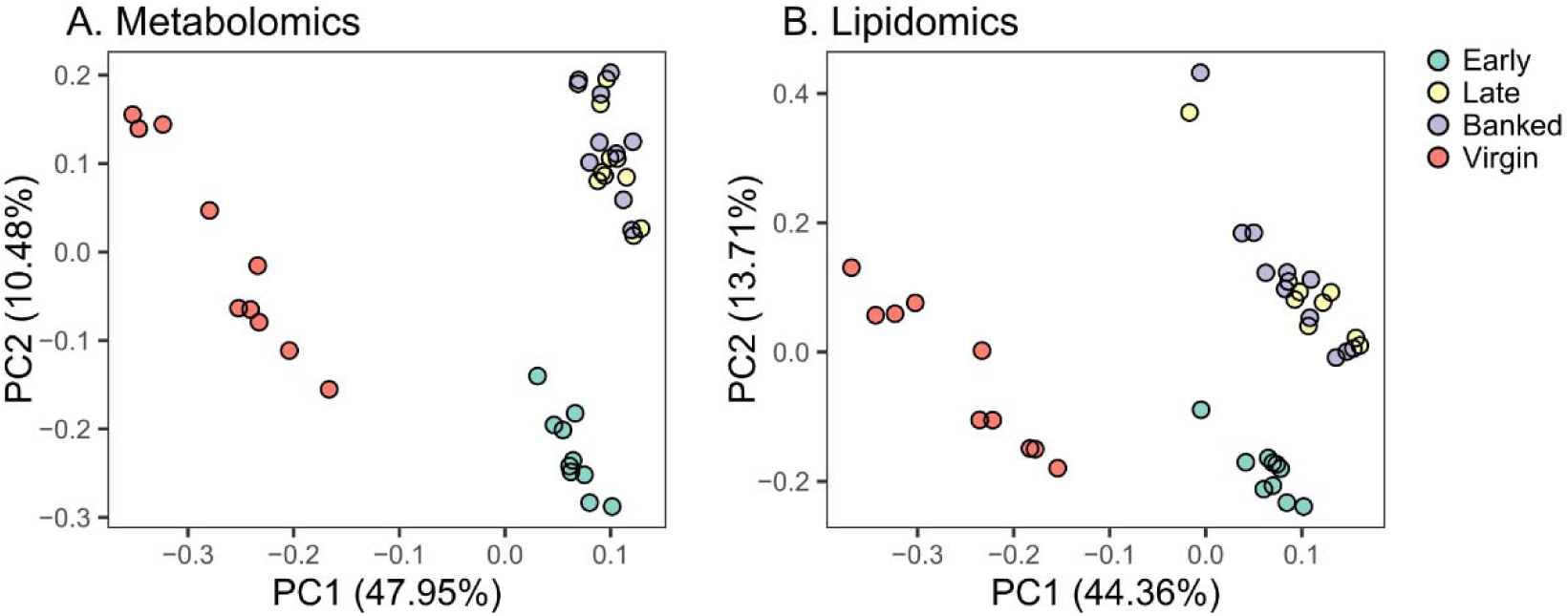
Untargeted metabolomics and lipidomics samples cluster strongly by group. In both sample types, compounds from positive and negative ion mode were pooled ahead of clustering. A) Metabolomics analysis relatively quantified 3,023 unique features, 238 of which were annotated, respectively. B) Lipidomics analysis relatively quantified 1,739 unique features, 258 of which were annotated.

Differential abundance testing shows that, of the 3,023 metabolite features and 1,739 lipid features relatively quantified, 1,022 and 525 were differentially abundant, respectively, in late versus early queens (**Supplementary Data 1-4**; 4% global FDR, Benjamini-Hochberg method). Only 86 and 109 of these features are confidently annotated, with patterns of abundance shown in **Figure 6**. By contrast, comparing banked and late queens (which were the same age and differed only in their level of ovary activation), we found only 378 differentially abundant metabolite features (32 of which were annotated) and 419 lipid features (73 of which were annotated). Among mated queens, six annotated compounds correlated with ovary mass, with one tentatively annotated as 4-hydroxy-3-methoxycinnamic acid (also known as ferulic acid) being the only one positively associated (**Figure S5**). It is worth noting that ferulic acid, a chemical compound commonly present in plant cell walls, has four possible isomers, including ferulic and isoferulic acid. Each isomer can exist as *cis* or *trans* stereoisomers. Although these compounds are not typically found in insects, insects may consume ferulic acid as part of their diet. Metabolomics and lipidomics profiles of all annotated compounds in all sample groups (including virgins) were also evaluated and shown in **Supplementary Figure S6 & S7,** with underlying data and statistical results available in **Supplementary Data 1-4**.

**Figure 6.**
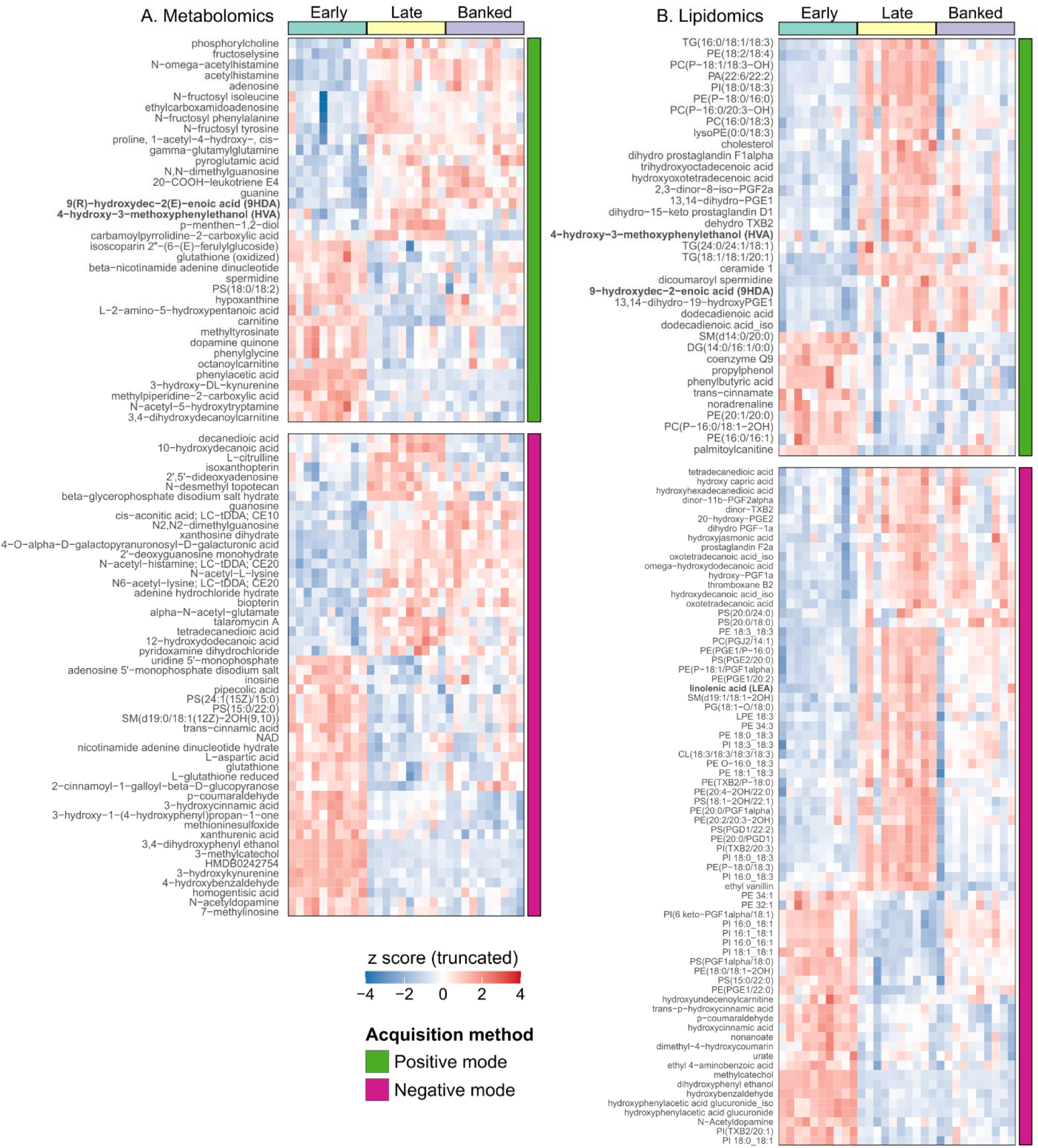
Metabolites and lipids extracted from heads vary by age among mated queens. Only annotated compounds are shown. False discoveries comparing late to early queens were controlled to 1% using the Benjamini-Hochberg method (4% additive global FDR, as four “families” are shown here). Bold labels represent queen retinue pheromone components. Detailed information for each compound annotation is available in **Supplementary Data 1-4** and a heatmap displaying all annotated compounds and all groups is in **Supplementary Figures S4 & S5.** A) Metabolomics: 86 annotated compounds had differential abundances in the late-early queen contrast. B) Lipidomics: 109 annotated compounds had differential abundances in the late-early queen contrast.

Comparing late and early queens, the most strongly differentially abundant (largest |fold change| and adjusted p < 0.01) annotated metabolites were N-acetyl-histamine, 2’,5’-dideoxyadenosine, beta-glycerophosphate, acetylhistamine, and N-omega-acetylhistamine, which are higher in late queens, and phenylacetic acid, 3,4-dihydroxyphenyl ethanol, 4-hydroxybenzaldehyde, 3-methylcatechol, and catalpol, which are higher in early queens. Among lipids, the most strongly differentially abundant compounds were dicoumaroyl spermidine, trihydroxyoctadecenoic acid, dihydro prostaglandin F-1α, hydroxy prostaglandin F-1α, and hydroxyhexadecanedioic acid (higher in late queens) and hydroxybenzaldehyde, dihydroxyphenyl ethanol, hydroxyphenylacetic acid glucuronide, methylcatechol, and hydroxyundecenoylcarnitine (higher in early queens). Many more differentially abundant features were identified (described in supplemental data) and most do not have roles previously described in honey bees.

Comparing late and banked queens, we found that the most strongly differentially abundant metabolites were carnitine and acetyl-L-carnitine (higher in banked queens) and citrulline, 2-hydroxy-3-methylpentanoic acid, 5-hydroxyindoleacetic acid, 4-Hydroxy-3-methoxycinnamic acid, and carbamoylpyrrolidine-2-carboxylic acid (higher in late queens). Among lipids, palmitoyl carnitine, hydroxybenzaldehyde, dihydroxyphenyl ethanol, isoforms of hydroxyphenylacetic acid glucuronide, and methyl catechol were higher in banked queens, whereas dodecenoylcarnitine, hydroxy-prostaglandin F1α, thromboxane B2, and hydroxydecanoic acid were higher in late queens.

In a separate analysis that included the virgin and banked queen samples, we found that amino acids, their derivatives, biogenic amines, prostaglandins, and thromboxanes were among the most strongly differentially regulated compounds (based on variable importance in projection (VIP) scores; **Figure 7**). In the metabolomics samples, citrulline was the most strongly differentially abundant compound, with the well-known neurotransmitters serotonin and octopamine also appearing among the top 15 differentially regulated compounds. With the exception of 2-ethyl suberic acid and suberic acid, levels were generally high in virgins and low in mated queens. The top compounds identified in the lipidomics samples were dominated by thromboxanes and prostaglandins, with generally low levels in virgins and higher levels in mated queens.

**Figure 7.**
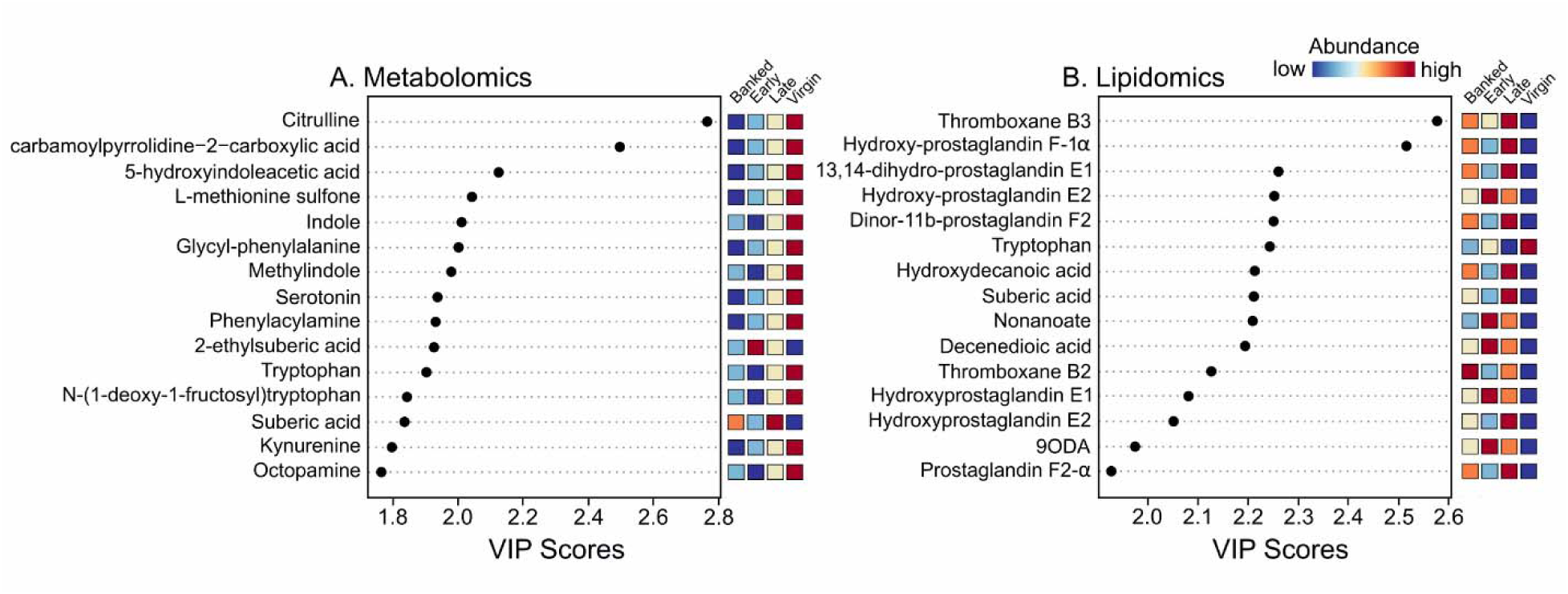
Variable importance in projection (VIP) scores for the top 15 metabolites and lipids extracted from queen heads. A VIP plot generated from the PLS-DA models ranked metabolites for their power to discriminate across groups. The higher the VIP scores for the metabolites, the more significant the contribution in the differences between groups. The relative abundance of metabolites is indicated by a colored scale from blue to red, representing the low and high, respectively. Top metabolites were generally at higher levels in the virgin group. In contrast mated queens which had remained in their mating nucs (late group) present higher levels of prostaglandins and thromboxanes.

## Discussion

Previous research by Rhodes *et al.*^21,29^, who tested short-term and sustained likelihood of acceptance and pheromone profiles in relation to queen age, in part motivated this work. Our data are largely in agreement; although the differences in acceptance that we observed between young and old queens were not significant, the trend for older queens to have higher acceptance rates is consistent with their observations. Our experimental design had three notable differences: 1) all queens in the introduction cohorts were confirmed to have been mated and laying eggs before introduction to recipient colonies (which was not the case in Rhodes *et al.*^21^); 2) we included an age-matched banked queen cohort to test the effect of laying status (and, by extension, ovary mass) on acceptance likelihood, and 3) we analyzed not only pheromone molecules but also a suite of other metabolites and lipids to gain insight into other changes in molecular profiles early in a queen’s life. These changes allowed us to deepen conclusions previously made by Rhodes *et al.*^21,29^ while also revisiting the “honest signal hypothesis”, on which there has been debate in the literature^19,31,55^.

The honest fertility signal hypothesis posits that workers can derive information about a queen’s fertility from pheromonal signals, which may then affect worker preference by allowing them to act in their own genetic best interest^16^. The mechanisms underlying this hypothesis have not been resolved, but our experiments show that if an honest signal does indeed exist, ovary activation is apparently not one of the fertility metrics conveyed. Previous studies suggested it could be^17,18^, but confounding factors of queen age and insemination quality render both studies effectively inconclusive. In light of our experiment showing that banked queens had just as high introduction success as age-matched free-range queens, if not higher (**Figure 3A & B**), despite having significantly smaller ovaries (**Figure 3C**), it is clear that ovary activation is not a defining indicator of acceptance likelihood. It is unlikely, therefore, that workers are perceiving chemical indications of ovary mass, *per se*, as being more attractive, as opposed to features which might normally covary with ovary mass (such as age, mating status, or absence of infections^56^).

Ovary mass is a measure of instantaneous fecundity, but it is the only fertility metric that is reversible (as opposed to ovariole number, sperm viability, or sperm count, which are either fixed or can only decline) and it stands to reason that an honest signal must be derived from an honest metric. We speculate that previously identified relationships between attractiveness and ovary mass may be the product of a chicken-or-egg scenario, whereby queens may actually develop larger ovaries due to improved attendance and feeding frequency, rather than larger ovaries causing higher attractiveness. Despite ambiguity in the causal direction, we can infer that honest pheromonal fertility signals should vary independently of ovary mass.

Our data and previously published data^19^ cumulatively point to 9-HDA as a candidate fertility signal. In our experiments, we found that 9-HDA varies independently of ovary mass among mated queens (**Figure 4C**) and positively associates with the queen cohorts that yielded the highest introduction success (**Figure 4A**). Although the fertility *potential* in queens belonging to the early cohort should not, in theory, differ from the late or banked cohorts, we speculate that they may be perceived as less attractive because the physiological changes induced by insemination^57–60^ have had little time to occur. Interestingly, Strauss *et al.* found that 9-HDA also tends to be more abundant in worker-laying queens compared to age-matched drone-laying queens^19^ ― a fertility characteristic that is not reversible. This compound thus deserves further attention as a candidate fertility signal and possible promoter of queen acceptance.

Similarly, we also found that HVA was more abundant in our high-acceptance cohorts and Strauss *et al.* found the same pattern of variation for HVA as for 9-HDA^19^. However, we found that HVA does not vary independently of ovary mass (**Figure 4B**) and is thus not likely a reliable fertility signal, though it may still be a determinant of worker preference. HVA is particularly interesting as a possible acceptance lubricant because it elicits a comparable response as dopamine in the worker brains^12^. While QMP, as a bouquet, suppresses worker dopamine signalling through downregulation of dopamine receptors, HVA alone appears to interact with those receptors to cause dopamine-like effect in the mushroom body^12^.

Foragers express high levels of dopamine receptors relative to nurses, and suppression of dopamine signalling reduces physical activity levels^12^. Therefore, HVA might be perceived as a reward, while contact with other QMP components reduces dopamine signalling to suppress worker activity and balling behavior. Indeed, Robinson *et al.* wrote that, in observation hives, most balling workers were non-aggressive, and both aggressive and non-aggressive balling workers tended to be older (> 12 d post-emergence), eventually becoming conditioned to foreign queens several hours after contact^61^. These observations could have been the real-time product of the interplaying dopamine signalling and dopamine suppressing effects of QMP.

Our ability to reliably detect HVA even among virgin samples highlights LC-MS/MS as an underutilized analytical tool for pheromone detection. This is notable because HVA abundance in virgin queens is sufficiently low for this compound to frequently fall below the limit of detection using conventional GC-MS systems^62–64^. Using an uncomplicated two-phase lipid and metabolite extraction protocol^13^ combined with LC-MS/MS, we assessed not only HVA in virgins but also hundreds of other metabolites and lipids, many of which significantly varied by queen cohort. Dopamine was significantly more abundant in virgin queens compared to mated queens (**Figure 7 & Figure S6**), consistent with past research^65,66^; however, other biogenic amines (*e.g.* serotonin, octopamine, and spermidine) were also more abundant in virgins and, to the best of our knowledge, have not yet been investigated in the context of queen development. Spermidine was also differentially abundant among the mated queen groups (**Figure 6A**), with higher levels in early compared to late cohorts. Spermidine treatment is known to increase vitellogenin expression in workers^67^, but perhaps it has other important functions early in a queen’s life.

While we had success with relative quantification of HVA and the analytical method generally performed well, we were not able to perform relative quantification all QRP components. Alcohols are generally difficult to observe by LC-MS/MS, which influenced our ability to evaluate PA, as its abundance in samples fell below our limit of detection (as determined using the PA standard). PA is therefore still best analyzed by GC-MS. In addition, 9-HDA appears in QRP as both R and S enantiomers, with 9(R)-HDA being the dominant isoform; however, the only commercially available chemical standard we could source was supplied as an 85%/15% mix of R/S entantiomers, and only 9(R)-HDA was visible in the chromatogram. While we are confident that we can distinguish 9(R)-HDA from 9(S)-HDA in our samples (**Figure S1**), we only report relative quantities of the former due to the poor performance of the standard. Finally, since the raw material for sample extraction consisted of whole heads, this means that some compounds may actually be derived from food leftover in the queen’s mouthparts. Among others, 10-HDA stands out in particular as being a potential food-derived compound, as it is a major component of royal jelly and there is severe disagreement in the literature around its relative abundance in virgin and mated queens^31^. Young virgins’ first meal as adults is often the leftover royal jelly in the cell from which they emerge; therefore, mouthpart contamination in heads or dissected mandibular glands could give rise to divergent quantities.

Our combined approach of analyzing pheromones, metabolites, and lipids from the same samples provided complementary data with no additional sample processing labor. While some QRP components (HVA, 9-ODA, and 9-HDA) and the non-QRP component 10-HDA were relatively quantified in both sample types, HOB and CA were present in metabolomics samples and LEA and MO were present in lipidomics samples. Both metabolomics and lipidomics samples clustered strongly according to queen age, however, with banked and age-matched free-range cohorts indistinguishable on PCA plots, suggesting that age and not laying status is the primary determinant of metabolic and pheromonal changes. Indeed, we identified far more metabolites and lipids which were differentially abundant between late and early cohorts (1,547 features) than between late and banked cohorts (797 features). Those compounds which were differentially abundant between late and banked queens, such as carnitine, citrulline, some prostaglandins, and many others, are likely involved in ovary activation and the associated metabolic changes.

Our lipidomics analysis including virgin samples shows that prostaglandins dominate the lipidomic changes, with generally low levels in virgins (**Figure 6B**), which, combined with some prostaglandins being positively associated with late (laying) vs. banked (not laying) queens, supports the notion that they are important for oogenesis or oviposition. Honey bees do have prostaglandin receptors and prostaglandin-associated enzymes^68–70^, and prostaglandins are positively linked to oogenesis and immunity in insects^71^, but the role of prostaglandins in honey bees has not yet been investigated. Interestingly, prostaglandins release oviposition behavior in crickets^34^, which is consistent with our observation that banked queens (which are not actively laying eggs) have lower levels of prostaglandins than late queens (which are constantly ovipositing). Similarly, we found that several thromboxanes, which are structurally similar to prostaglandins, were upregulated in mated queens over virgins (**Figure 7 & Figure S7**). In the Lepidopteran *Spodoptera exigua*, thromboxanes are immunostimulatory^72,73^, and a thromboxane receptor has been characterized in *Aedes* and *Anopheles* mosquitoes^74^, but neither thromboxanes nor their receptors have been investigated in honey bees.

We anticipate that the data presented in this manuscript will serve as a resource to guide further metabolite and lipid inquiries in honey bees. However, we note that only the QRP components, 10-HDA and L1 annotations have been unequivocally confirmed with analytical standards. High-confidence identifications (L2 annotations, as indicated in **Supplementary Data 1-4**) are considered reliable, but assignments should be confirmed experimentally for complete accuracy. To facilitate this effort, spectral data are openly accessible for additional interrogation or improvements in annotation as new pathways are uncovered and new standards become accessible. These comprehensive spectral data will serve as a legacy dataset for ongoing exploration.

## Conclusion

We present a map of metabolic, lipidomic, and pheromonal transitions that occur during the early stages of a queen’s life. We used these data to investigate patterns underlying age-related trends in queen acceptance that are independent of ovary mass, bringing the relevance of ovary mass as an honest fertility metric into question. Likelihood of sustained queen acceptance increases with queen age from 76% for 10-12 d old queens to 90% for one month old banked queens. This difference was marginally non-significant (p = 0.070) but the trend is consistent with previous research documenting significant but otherwise similar patterns^21^. Queen banking for 18 days did not affect acceptance rates or retinue pheromone profiles, but pheromones, metabolites, and lipids profoundly varied by age. We hope this rich dataset and uncomplicated methods will provide a path forward for a new area of research integrating pheromonal profiles and metabolism in honey bees.

## Acknowledgements

We would like to thank The Scandia Honey Company for generously supporting this research, and Intko Supply Ltd for providing us with 9-ODA and 9-HDA analytical standards. We also thank J. Kearns, D. Baker, M. McKay, L. Holmes, and J. Todoschuk for assistance in the field.

## Author contributions

SEH conceptualized the study and grants to SEH and LJF funded the work. SEH conducted all field work associated with this study in collaboration with personnel at Scandia Honey Company. AM wrote the manuscript with editing assistance from SEH, LJF, and AC. AM prepared the samples, analyzed the data, and produced the figures, with assistance from AC. AC optimized LC-MSMS methods, acquired all mass spectrometry data, and curated the data according to MSI standards.

