## Supplementary Figures S1-S7 for "Parallel pheromone, metabolite, and lipid analyses reveal patterns associated with early life transitions and ovary activation in honey bee (*Apis mellifera*) queens"

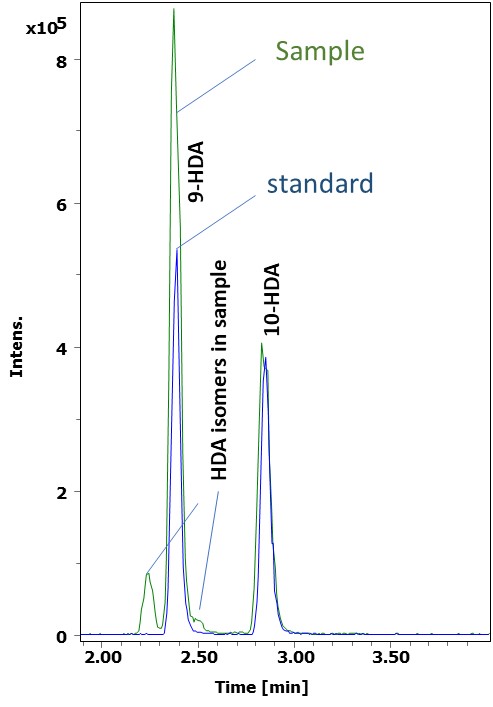

**Figure S1. 9-HDA and 10-HDA pheromone standards versus example sample chromatogram.** The 9-HDA standard was commercially available only as a mixed product of 85% R and 15% S enantiomer proportions. Only the R enantiomer was observed in the mixed standard; however, the low-abundance isomer to the left of the major 9-HDA peak, which is observable in queen samples, is very likely the S enantiomer.

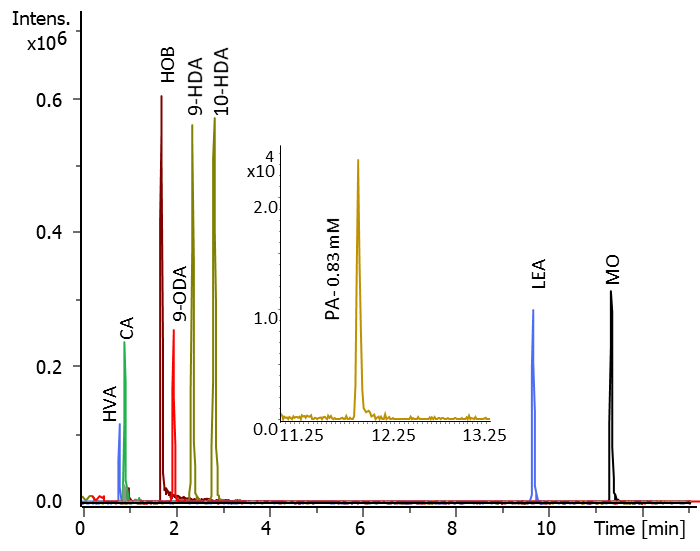

**Figure S2. Example of chromatograms of components of QRP (standards).** Data were obtained from a 1 µg/mL injection of 1 µl. Note that 9-HDA refers to the R enantiomer only.

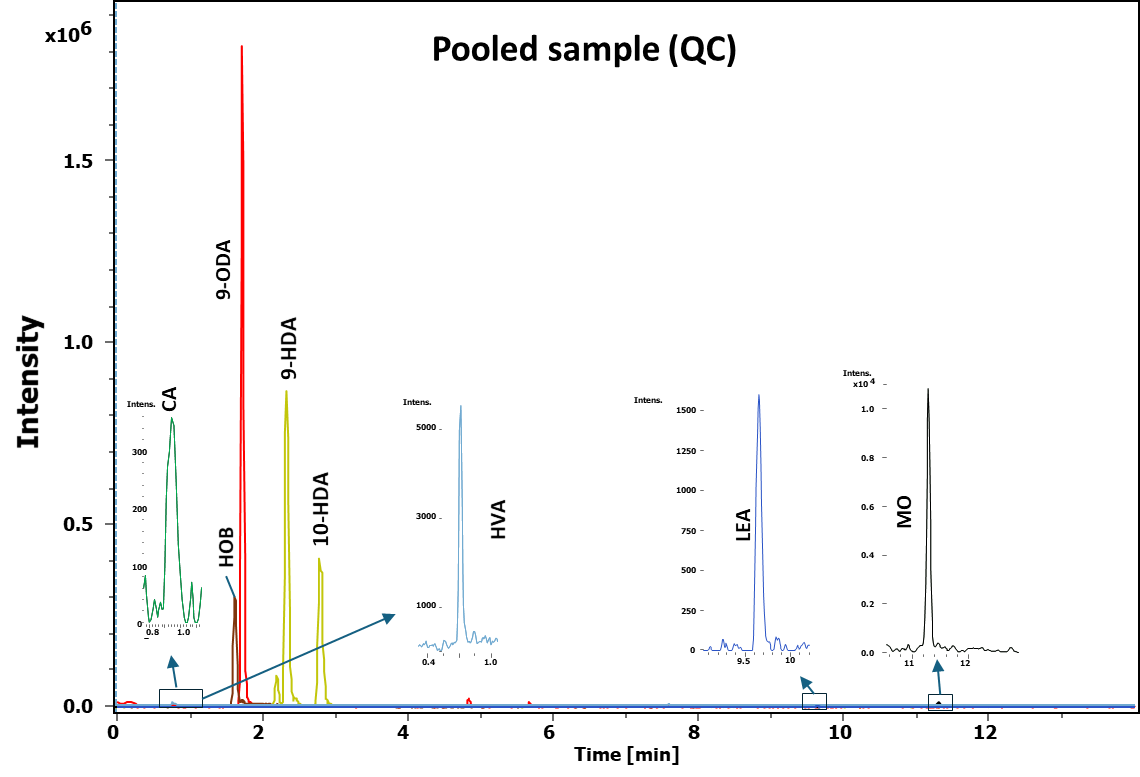

**Figure S3. Extracted ion chromatogram (XIC) of QRP compounds detected in a pooled sample (lipidomics fraction).** The XIC was as follows. ESI-, HOB, 151.0399; 9-ODA, 183.1026; LEA, 277.2134. ESI+, CA, 131.0494 (fragment); 9- and 10-HDA, 187.1329, HVA 191.0676 (sodium adduct); MO, 297.2785. Retention time matched with standards.

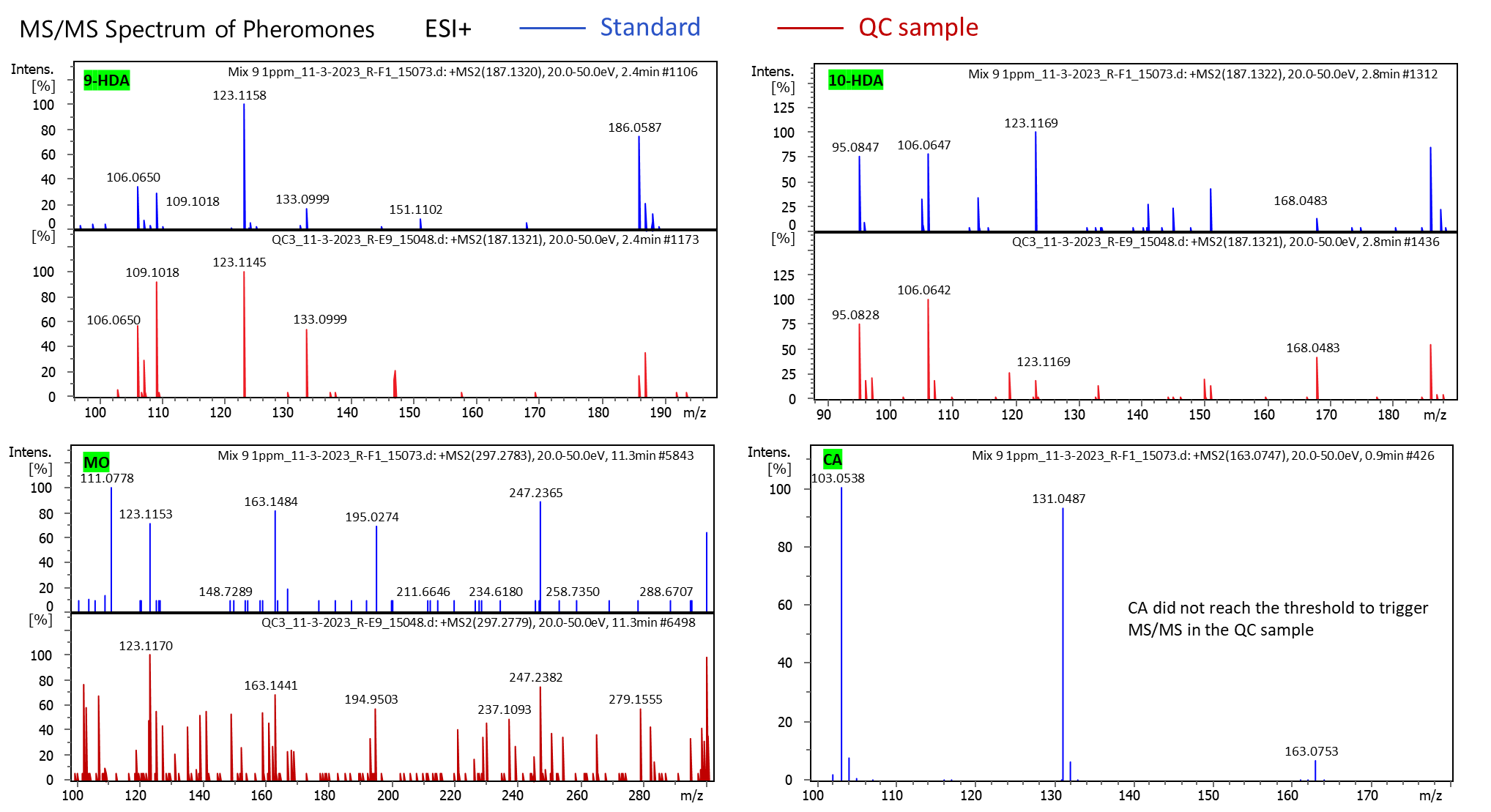

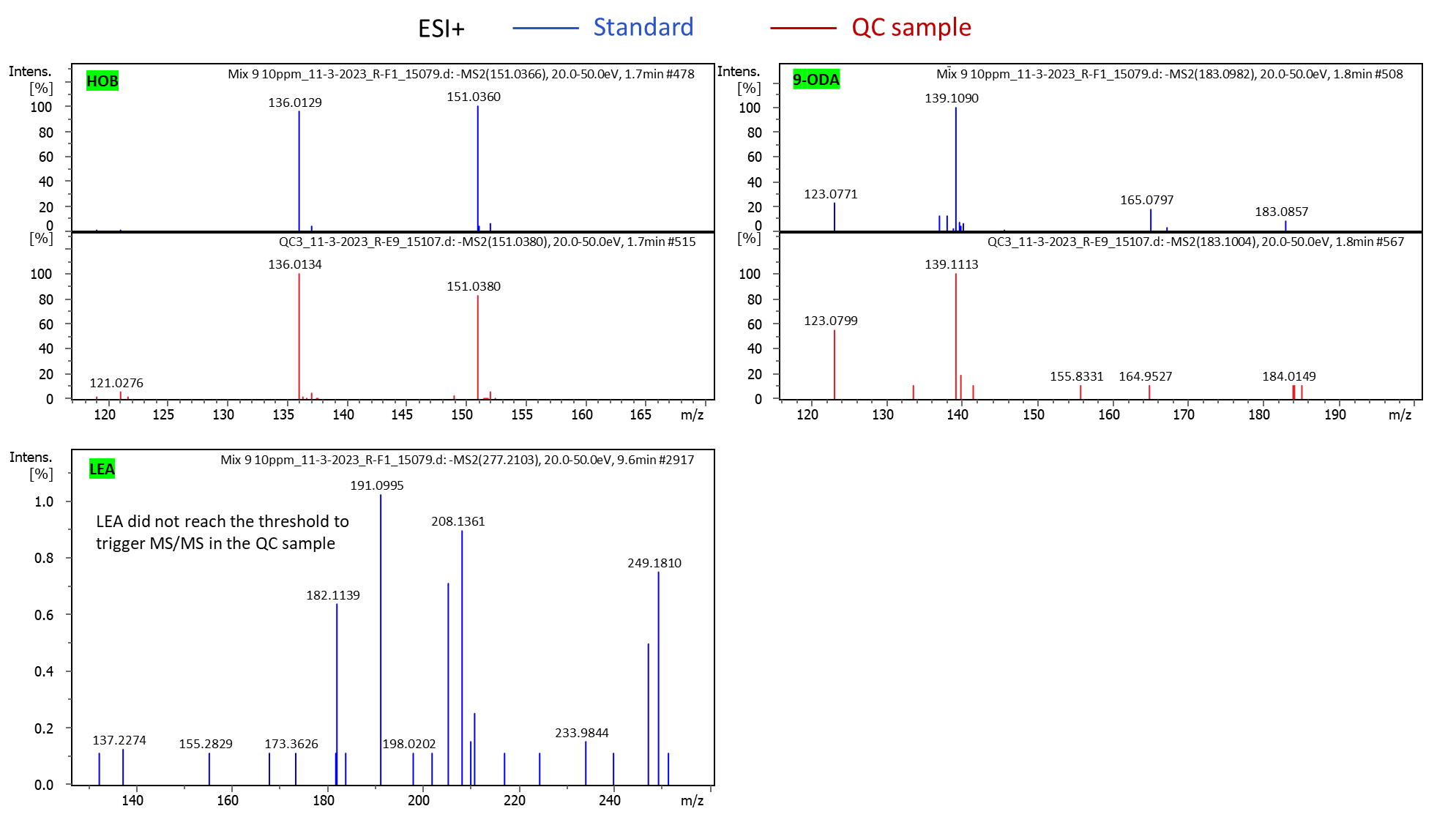

**Figure S4. Example MS/MS spectra of QRP standards and samples.** CA and LEA did not reach a sufficient intensity to trigger MS/MS spectra acquisition.

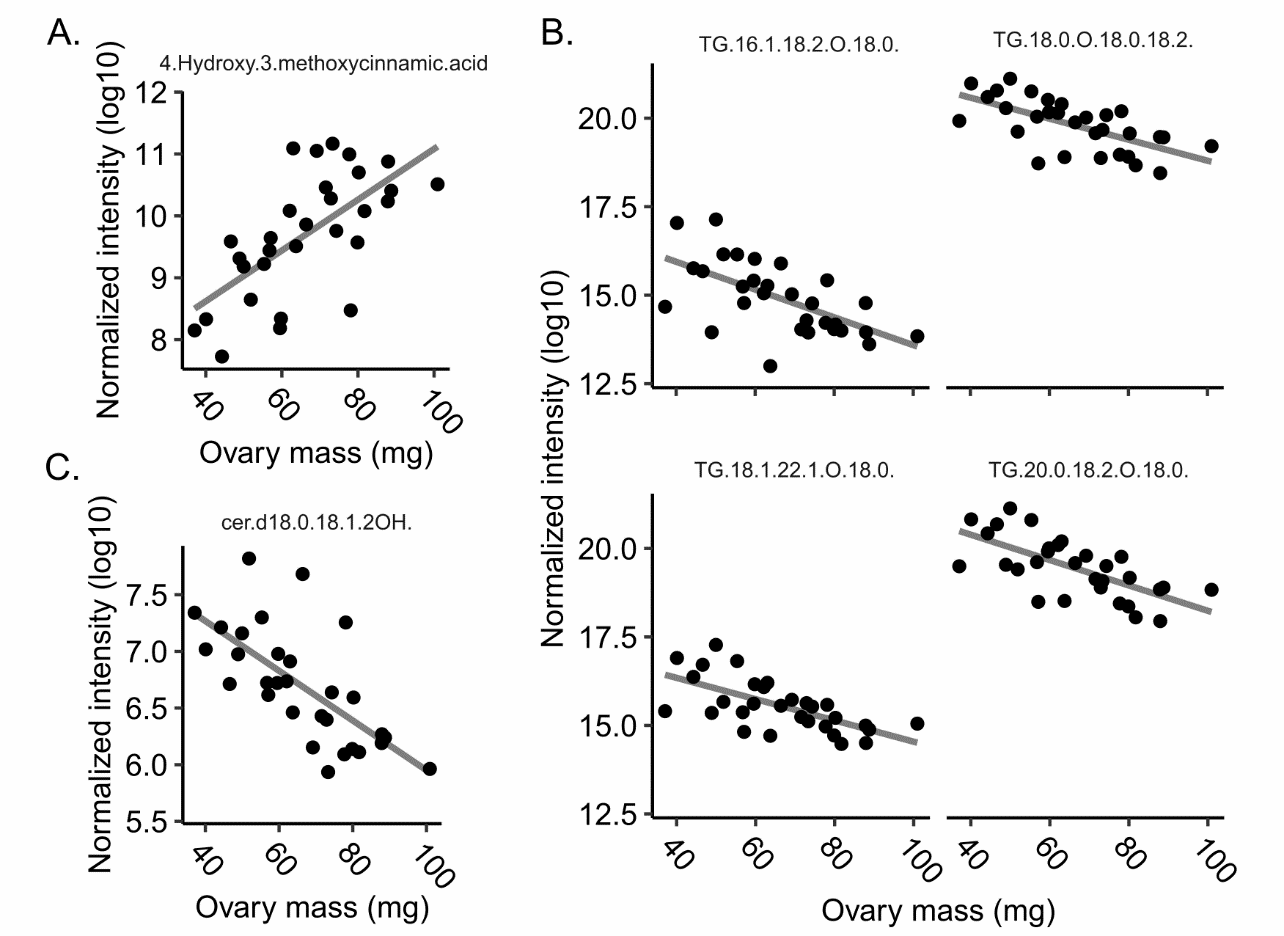

**Figure S5. Relationships between compound abundance and ovary mass.** Virgin queen samples were excluded and only identified compounds are shown. A total of 28 and 39 compounds were significantly linked to ovary mass in metabolomics and lipidomics samples, respectively. False discovery rates were controlled at 1% (Benjamini-Hochberg method). A) Metabolomics, negative ion mode. B) Lipidomics, positive ion mode. C) Lipidomics, negative ion mode.

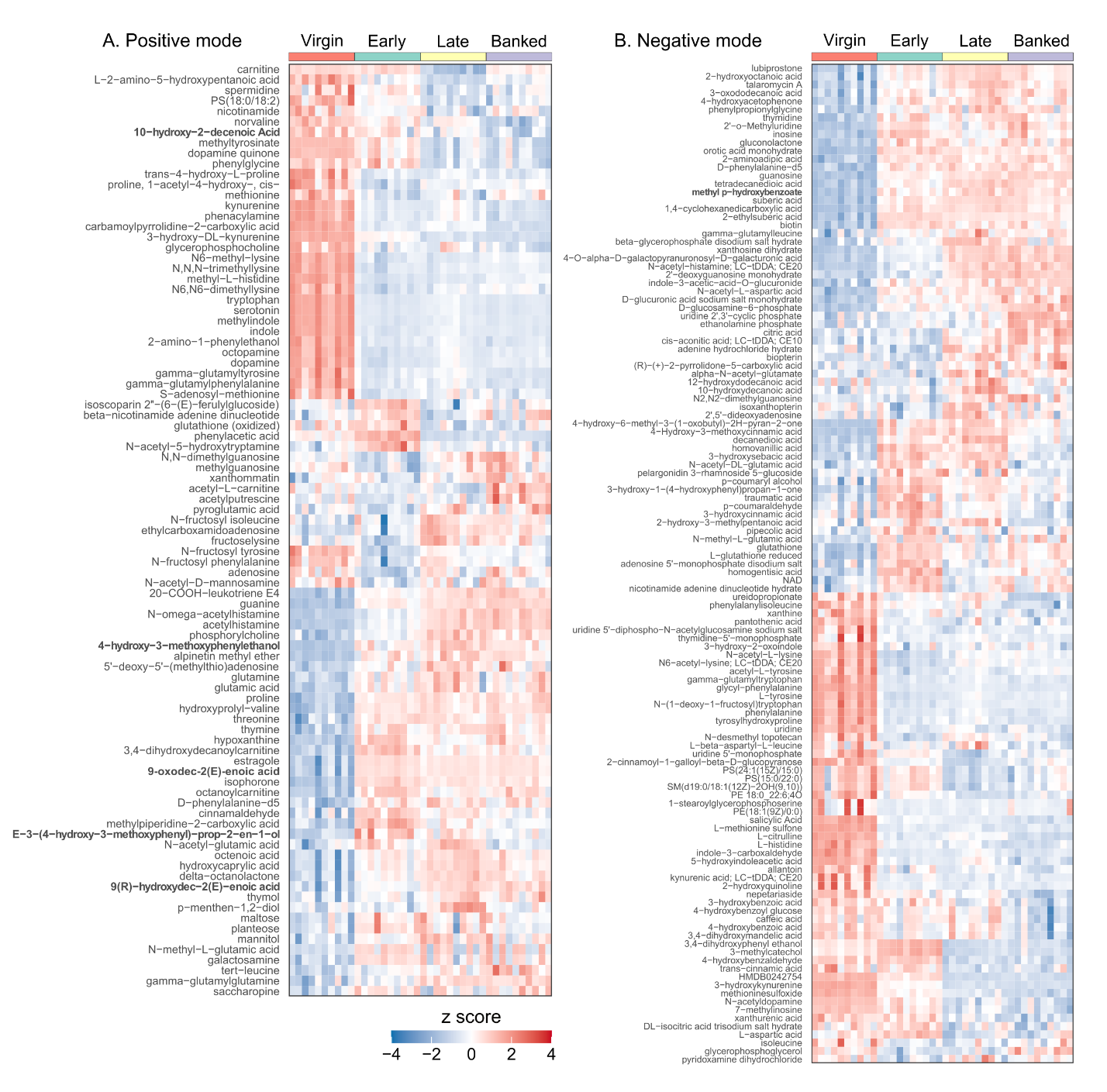

**Figure S6. All annotated compounds in metabolomics samples**. Compounds without names/identities are not shown.

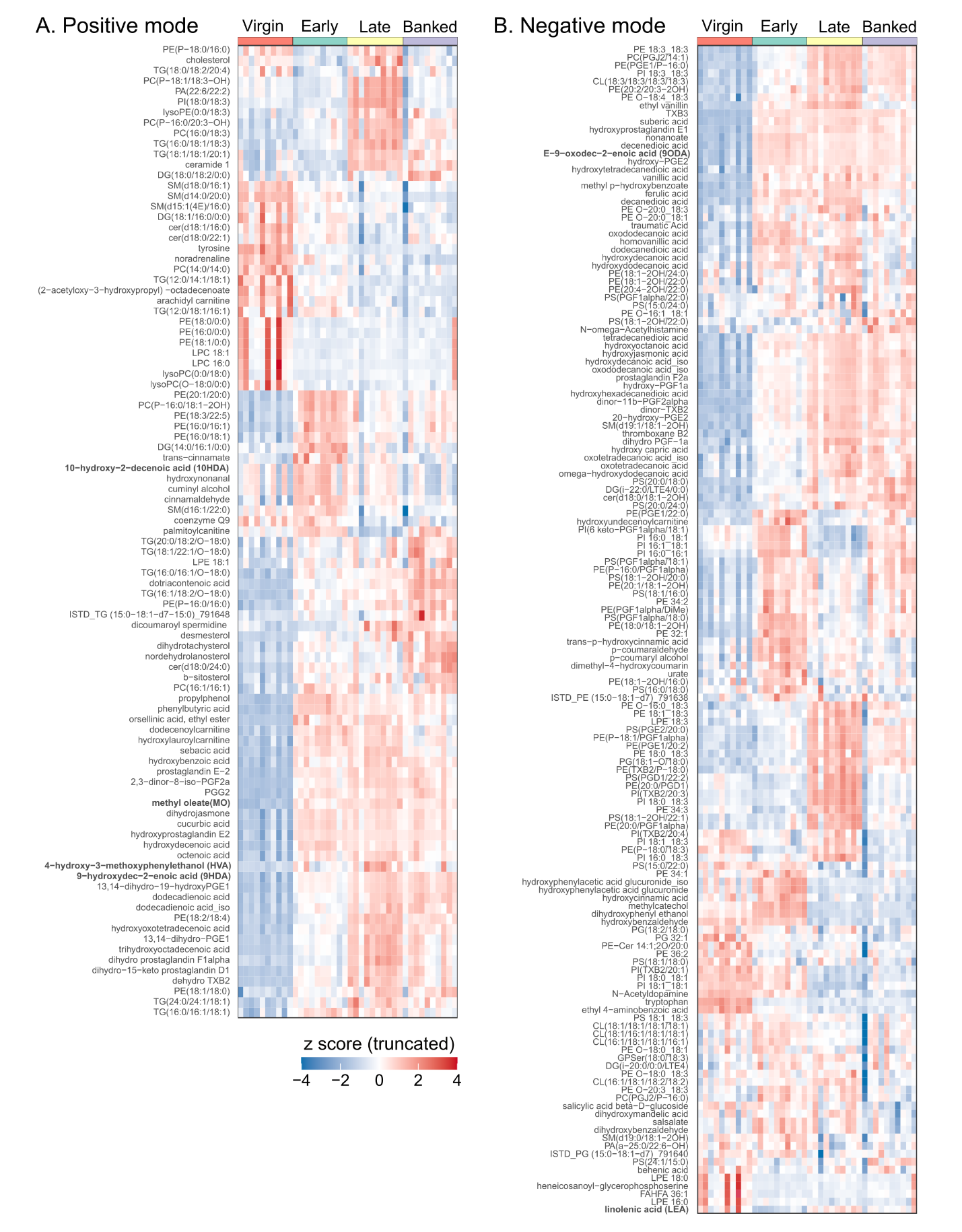

**Figure S7. All annotated compounds in lipidomics samples.** Compounds without names/identities are not shown.
