## Supplementary File 1 for "Parallel pheromone, metabolite, and lipid analyses reveal patterns associated with early life transitions and ovary activation in honey bee (*Apis mellifera*) queens"

### Queen pheromone, metabolite, and lipid extraction for LC-MS/MS

#### Introduction

This protocol uses a dual extraction approach based on Chen *et al.* (2013) to sample lipids, metabolites, and pheromones all from a single queen head. All reagents should be of analytical grade for LC-MS/MS.

Abbreviations:
BHT: Butylated hydroxytoluene
MTBE: Methylated tert-butyl ether
CUDA: 12-[[(cyclohexylamino)carbonyl]amino]-dodecanoic acid
ppm: Parts per million
QC: Quality control

#### Materials

- BHT Methanol (0.01% w/v)
  - Make a 1% BHT methanol stock (100 mg in 10 mL)
  - Dilute stock 1:100 in methanol to make working solution
  - Store working solution in freezer (-20 °C)
- 75% Methanol:Water (v/v) extraction solvent using 0.01% BHT methanol
  - Add CUDA internal standard to a final concentration of 1 ppm
  - Add methionine-d3, caffeine-$^{13}C_3$, and ferulic acid d3 to final concentrations of 1 ppm each
  - Add SPLASH LIPIDOMIX (Avanti Polar Lipids) internal standard mixture to a final concentration of 5 ppm
  - Note: Make solution fresh or store short-term in freezer (-20 °C)
  - Note: Normally an 80% methanol:water solution would be used, but queen heads contain a substantial amount of water, so the water component is reduced accordingly here
- MTBE extraction solvent (neat)
  - Store neat solvent in freezer for easier pipetting
- 50% Methanol:Water (v/v) metabolite resuspension solvent
- 70% Acetonitrile:Isopropanol (v/v) lipid resuspension solvent
- 80% Methanol:Water (v/v) solvent for preparing queen pheromone standards
- Pheromone standards
  - See Table 1 below for more information on the queen pheromone standards
- Homogenization tubes (*e.g.* Tough tubes from Ulti Dent Scientific, 2641-0B)
- Ceramic beads (2.8 mm diameter)
- Tissue homogenizer (*e.g.* Precellys-24, Bertin Instruments)

- Refrigerated microcentrifuge
- Thermomixer
- Vortexer
- Other laboratory materials
  - micropipets
  - pipet tips, colorless
  - 1.5 and 2.0 ml microfuge tubes, colorless
  - LC-MS/MS autosampler vials

#### Procedure

##### Homogenization and extraction

1. Prepare pheromone standards (Table 1) at 1 µg/ml in 80% BHT methanol

**Table 1. Queen pheromones**

| | A | B | C | D |
|---|---|---|---|---|
| 1 | Name | Short name | Monoisotopic mass | Supplier |
| 2 | E-9-oxodec-2-enoic acid | 9-ODA | 184.1099 | Intko Supply Ltd, Chilliwack, BC, Canada |
| 3 | 9(R)-hydroxydec-2(E)-enoic acid | 9(R)-HDA | 186.1256 | Intko Supply Ltd, Chilliwack, BC, Canada |
| 4 | 9(S)-hydroxydec-2(E)-enoic acid | 9(R)-HDA | 186.1256 | Intko Supply Ltd, Chilliwack, BC, Canada |
| 5 | methyl p-hydroxybenzoate | HOB | 152.0473 | MilliporeSigma, Burmington, MA, USA |
| 6 | 4-hydroxy-3-methoxyphenylethanol | HVA | 168.0786 | MilliporeSigma, Burmington, MA, USA |
| 7 | methyl oleate | MO | 296.2715 | Cayman Chemical, Ann Arbor, MI, USA |
| 8 | E-3-(4-hydroxy-3-methoxyphenyl)-prop-2-en-1-ol | CA | 180.0786 | Cayman Chemical, Ann Arbor, MI, USA |
| 9 | linolenic acid | LEA | 278.2246 | Cayman Chemical, Ann Arbor, MI, USA |
| 10 | 10-Hydroxy-2(E)-decanoic acid | 10-HDA | 186.1256 | Cayman Chemical, Ann Arbor, MI, USA |

2. Excise whole queen heads and deposit in a homogenization tube with 4 ceramic beads (2.8 mm diameter)

   Note: Prepare a minimum of three blank samples in parallel, following identical steps but without a tissue sample added

   Note: Prepare samples in parallel wherever possible to minimize batch effects

3. Homogenize ~50 mg of sample (one head) in 400 µl methanol extraction solvent in a homogenizer (for example, a Qiagen TissueLyser or Precellys 24 tissue homogenizer). Homogenize at a frequency of 5,000 Hz for 30 s, then rest on ice for  min. Repeat 3x.

4. Add 1 mL ice-cold MTBE

5. Incubate samples for (minimum) 1 hour at room temperature, shaking 1000 rpm

6. Transfer 1400 µl to a new 2 mL clear tube

7. Centrifuge at 14,000 g for 10 minutes (4 °C) and transfer 1200 µl of supernatant to new tube

8. Add 214 ul water to induce phase separation, vortex briefly and incubate 10 min at room temperature

9. Centrifuge 15 min at 14 000 *g* (4 °C)

10. Transfer exactly 250 µl upper fraction (lipids) and 250 µl lower fraction (metabolites) to new tubes.

11. Speedvac metabolite fraction to dryness (2.5 h, room temperature) and store at -70 °C until resuspending. Store lipid fraction at -70 °C and speedvac only when ready to analyze to help prevent oxidation

#### LC-MS/MS analysis

12. Run pheromone standards ahead of samples to confirm detection and adequate separation of components.

    A 1 µl injection volume for standards at a concentration of 1 µg/ml should be sufficient, but choose the injection volumes and concentrations that are suitable for your LC-MS/MS system's needs.

    HOB and CA are best quantified in the metabolomics suspension whereas LEA, and MO are best quantified in the lipidomics suspension and 9-ODA, 9(R/S)-HDA, and HVA are quantified in both sample types.

13. Resuspend the metabolite fraction in 250 µl metabolite resuspension solvent. Centrifuge at 10,000 rcf (10 min, room temperature) to remove particulates and transfer 200 µl to an LC-MS autosampler vial

    Note: Pool 20 µl from each sample to produce a QC sample

    Note: It is good practice to inject QC and blank samples every 10 injections

    Note: It is essential to randomize sample injection orders between groups

    The exact LC settings may vary on different systems, but we have found the following parameters effective using an Impact™ II high-resolution mass spectrometer (Bruker Daltonics, Bremen, Germany) coupled with a Vanquish Horizon UHPLC system (Thermo) with an Inertsil Ph-3 UHPLC column (2 µm, 150 x 2.1 mm) (GL Sciences) equipped with a Ph-3 guard column (2 µm, 2.1 x 10 mm) (Table 2)

**Table 2. LC settings for metabolomics analysis**

| | A | B |
|---|---|---|
| 1 | Parameter | Description |
| 2 | Phase A | Water with 0.1% (v/v) formic acid |
| 3 | Phase B | Methanol with 0.1% (v/v) formic acid |
| 4 | LC gradient | 0 min (5% B), 0–1 min (5% B), 1–8 min (35% B), 8–10.5 min (99% B), 10.5–14 min (99% B), 14–14.5 min (5% B), and 14.5–18 min (5% B) |
| 5 | Column temperature | 55 °C |
| 6 | Autosampler temperature | 4 °C |
| 7 | Flow rate | 0.3 mL/min |
| 8 | Injection volume | 3 µl (for both positive and negative modes) |

14. Follow the same procedure to suspend lipid samples in lipid resuspension solvent, including QC, blank, and randomization procedures.

The exact LC and instrument settings may vary on different systems, but we have found the following parameters effective using an Impact$^{TM}$ II high-resolution mass spectrometer (Bruker Daltonics, Bremen, Germany) coupled with a Vanquish Horizon UHPLC system (Thermo) with an ACQUITY UPLC CSH C18 analytical column (130Å, 1.7 µm, 2.1 mm X 100 mm, Waters) (Table 3)

**Table 3. LC settings for lipidomics analysis**

| | A | B |
|---|---|---|
| 1 | Parameter | Description |
| 2 | Phase A | Water with 10 mM ammonium formate and 0.1% formic acid |
| 3 | Phase B | 10% acetonitrile and 90% isopropanol (v/v) with 10 mM ammonium formate and 0.1% (v/v) formic acid |
| 4 | LC gradient | 0 min (20% B), 0–2 min (20% B), 2–11 min (80% B), 11–11.5 min (99% B), 11.5–13.2 min (99% B), 13.2–14 min (5% B), and 14–18 min (5% B) |
| 5 | Column temperature | 65 °C |
| 6 | Autosampler temperature | 4 °C |
| 7 | Flow rate | 0.4 mL/min |
| 8 | Injection volume | 1 µl (positive mode) and 2 µl (negative mode) |
